# A lineage-specific nascent RNA assay unveils principles of gene regulation in tissue biology

**DOI:** 10.1101/2024.10.15.618417

**Authors:** Gopal Chovatiya, Alex B Wang, Philip Versluis, Chris K Bai, Sean Y Huang, Michael DeBerardine, Judhajeet Ray, Abdullah Ozer, John T Lis, Tudorita Tumbar

## Abstract

Gene regulatory mechanisms that modulate RNA Polymerase II activity are difficult to access in mammalian tissues composed of multiple cell lineages. Here, we develop a nascent RNA assay (PReCIS-seq) that measures lineage-specific transcriptionally-engaged Pol II on genes and DNA enhancer elements in intact mouse tissue. By employing keratinocytes as a prototype lineage, we unearth Pol II promoter-recruitment versus pause-release mechanisms operating in adult skin homeostasis. Moreover, we relate active enhancer proximity and transcription factor binding motifs on promoters to Pol II activity and promoter-proximal pausing level. Finally, we find Pol II firing rapidly into elongation on lineage identity genes and highly paused on cellular safeguarding genes in a context-dependent manner. Our work provides a basic platform to investigate mechanistic principles of gene regulation in individual lineages of complex mammalian tissues.

## Introduction

Mammalian tissue development and homeostasis implicate cell state transitions that are difficult to recapitulate *ex vivo*, outside the intact tissue milieu (*1*). Such transitions employ gene regulatory mechanisms controlling transcriptional activity at two major rate-limiting steps: i) initiation, with RNA Polymerase II (Pol II) recruitment to promoters, followed by rapid transition to promoter-proximal pausing (*2*) and ii) release of paused Pol II into productive elongation (*3, 4*). Pol II recruitment is modulated by pioneer transcription factors (*5*), whereas promoter-proximal Pol II pausing requires binding of the negative elongation factor (NELF) and DRB sensitivity-inducing factor (DSIF) (*3*). Pol II pause-release is driven by the positive transcription elongation factor-b (P-TEFb) complex, which phosphorylates Ser2 of Pol II c-terminal domain, as well as NELF and DSIF. This leads to dissociation of NELF from paused Pol II and conversion of DSIF to a positive elongation factor (*3*). Various mechanisms including sequence-specific transcription factors, co-regulators, chromatin structure, specific histone modifications, and other DNA elements such as enhancers and insulators act on specific gene promoters to regulate the level of Pol II pausing and pause-release (*6-9*). Importantly, various genetic, biochemical, and genomic approaches suggest that pause-release regulates cell signaling, cell cycle, and cell differentiation (*10-12*). Moreover, pausing and pause-release mechanisms may enhance gene induction speed, establish permissive chromatin, augment the synchronicity of gene induction in fields of cells during development, integrate regulatory signals, and act as a checkpoint for coupling RNA elongation and processing (*11, 12*).

Current methodologies for genome-wide Pol II activity mapping provide comprehensive insights into Pol II promoter-proximal dynamics and nascent transcription (*11, 13, 14*). However, employing these methodologies for specific cell lineages necessitates tissue dissociation and cell isolation (*15, 16*), which perturbs the natural cell-state and the tissue microenvironment essential for the biological transition in question (*17*). Previous elegant efforts for measuring mRNA levels in specific cell-types from intact mouse tissues capture a snapshot of transcriptional activity within hours of transcription (*18, 19*). However, these approaches cannot access transcriptionally-engaged Pol II, therefore the early steps of transcription activation remain hidden. Therefore, a lineage-specific, genome-wide assay for measuring transcriptionally-engaged Pol II activity in intact mammalian tissue is urgently needed.

Here, we develop PReCIS-seq (<u>P</u>recision <u>R</u>un-on in <u>C</u>ell-type-specific *<u>I</u>n vivo* <u>S</u>ystem followed by sequencing) which implements high resolution precision transcriptional run-on (e.g., PRO-seq) (*20*) to specific cell lineages in intact mouse tissue (Fig. 1A). To this end, we generated a knock-in mouse line which enables cell-lineage-specific, Cre-LoxP-driven replacement of the endogenous Pol II with a GFP-tagged Pol II. As a prototype cell linage, we employ keratinocytes– an essential skin building block with body protective functions (*21*) that maintains a distinct transcriptomic profile throughout organismal life (*22*). To obtain a global view of the keratinocyte transcriptome in intact skin tissue, we apply PReCIS-seq at three major developmental stages: 1) embryonic day (E)16.5 as mature skin epidermal structures are formed; 2) adult skin resting stage at postnatal day (PD) 21 (telogen); and 3) adult skin growth stage at PD24 (anagen), when keratinocytes increase proliferation preceding tissue expansion and differentiation (*23*). This enabled classification of developmentally controlled functional gene sets based on Pol II levels at the promoter-proximal pause region and gene body, resulting from Pol II recruitment and pause-release. Simultaneously, PReCIS-seq enabled genome-wide identification of transcriptionally-active enhancer candidates. Our platform uncovers mechanistic gene regulatory principles of mouse tissue biology enabling new access points into the molecular basis of human disease.

**Fig. 1.**
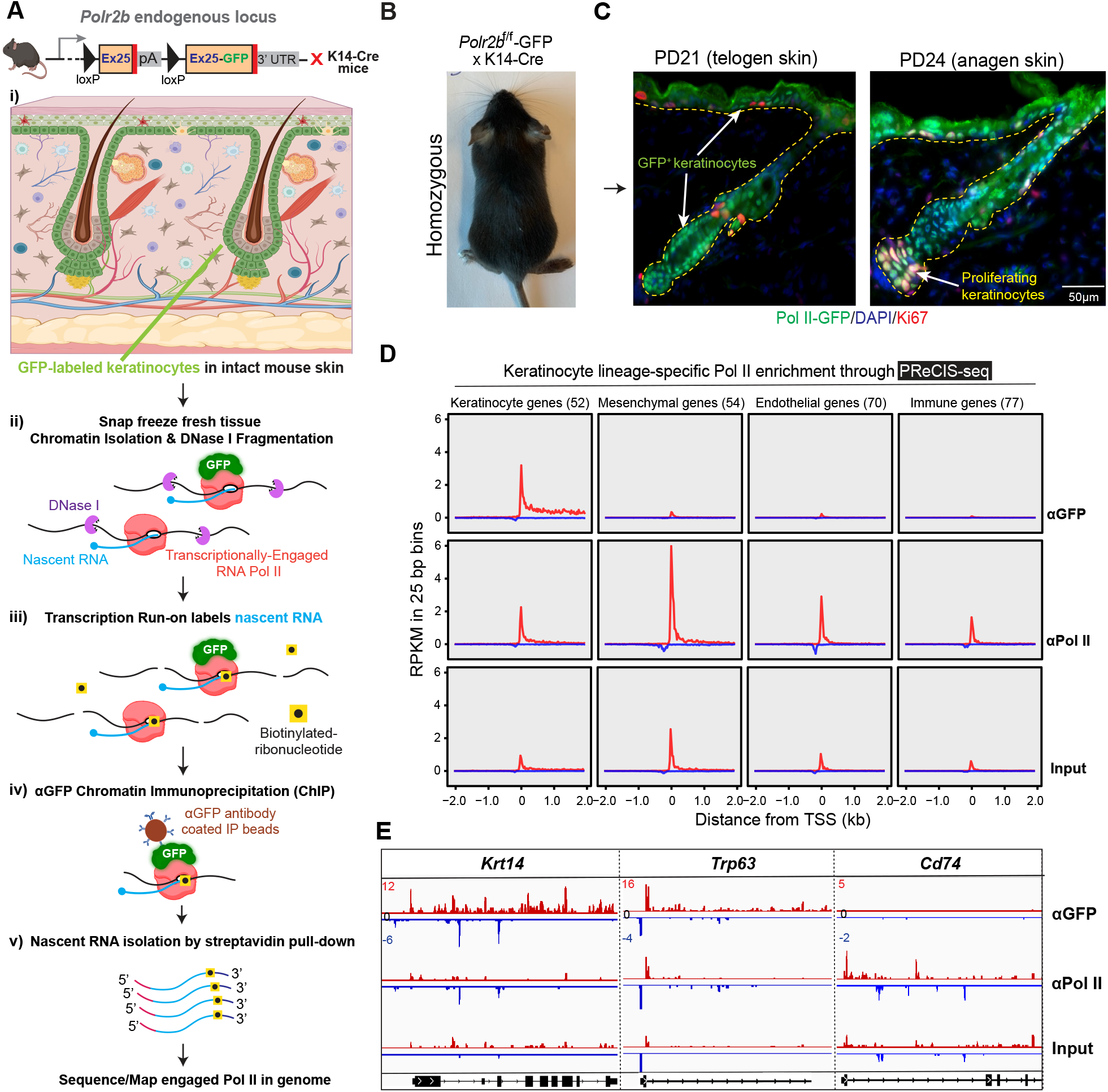
Precision Run-on in Cell-type-specific *In vivo* System followed by Sequencing (PReCIS-seq) maps transcriptionally-engaged RNA Pol II in intact mouse tissue. (**A**) Schematic depiction of PReCIS-seq assay (see also fig. S1A) in mouse back skin. (**B**) Homozygous *Polr2b*^f/f^-GFP transgenic mouse at postnatal day (PD) 21 (telogen) after crossing with mice bearing a keratinocyte-specific K14-Cre driver. (**C**) Immunofluorescence stainings of GFP and Ki67 proliferative cells and DNA DAPI staining in indicated colors in back skin sections from *Polr2b*^f/f^-GFP; K14-Cre mice. (**D**) Metaplots of PReCIS-seq and input data from telogen samples displaying density of Pol II around 2kb from TSS (mm39 RefSeq) on four major skin-lineage-specific gene sets. (**E**) Genome browser views of PReCIS-seq telogen data for *Krt14*, a keratinocyte-specific gene; *Trp63*, an epidermis differentiation transcription factor; and *Cd74*, a non-lineage, immune-cell-specific gene. The sense and anti-sense strands are shown on different scales and colors; red and blue, respectively. *Krt14* gene orientation is flipped for visualization purposes.

## Results

### PReCIS-seq reveals lineage-specific nascent transcriptomics in intact mouse tissue

Precise and highly-sensitive mapping of RNA Pol II occupancy and nascent transcription of specific cell lineages in their natural tissue environment, without cell isolation, is critical for rigorously identifying and assessing the role and mechanisms of transcription regulation in cell-type development, homeostasis and disease. Here, we engineer a mouse model where cell-type-specific Cre activity can fuse GFP to the 3’ end of the endogenous second largest subunit of RNA Pol II (*Polr2b*), resulting in the RPB2-GFP fusion protein (herein referred to as Pol II-GFP) (Fig. 1A and fig. S1, A and B; see Materials and Methods). In our first test of this method, the Keratin (K) 14-Cre driver excises the intervening sequence in the *Polr2b*^f/f^-GFP construct allowing production of a Pol II-GFP fusion protein exclusively in keratinocytes. The transcriptionally-engaged Pol II-GFP in a chromatin fraction can then be enriched by immuno-precipitation with anti-GFP (αGFP) (Fig. 1A_i_) (*24*). To preserve *in vivo* Pol II transcription engagement patterns, freshly collected whole tissue is immediately snap frozen, followed by chromatin isolation, DNA fragmentation, and transcription run-on with biotin-ribonucleotides to label nascent RNAs (Fig. 1A_ii-iii_). Next, αGFP chromatin immunoprecipitation (ChIP) pulls down complexes of Pol II-GFP and the associated nascent RNAs from Cre-targeted cells (Fig. 1A_iv_). Streptavidin pull-down of biotin-labeled nascent RNAs is followed by strand-specific RNA library generation, sequencing, and genome-wide mapping to identify precise locations of transcriptionally-engaged Pol II (Fig. 1A_v_). <u>P</u>recision <u>R</u>un-<u>O</u>n sequencing (PRO-seq) is a widely-used assay employed in cultured cells and bulk tissues or tumors to map transcriptionally-engaged Pol II with high sensitivity at single base-pair resolution (*20*). <u>P</u>recision <u>R</u>un-on in <u>C</u>ell-type-specific *In vivo* <u>S</u>ystem followed by sequencing (PReCIS-seq) integrates PRO-seq with αGFP immunoprecipitation of cell-type-specific Pol II-GFP enabling nascent transcription profiling of specific cell lineages in a complex tissue sample (Fig. 1A).

To create a Cre-inducible Pol II-GFP mice, we targeted the endogenous *Polr2b* gene locus in mouse embryonic stem cells (mESCs) generating the *Polr2b*^f/f^-GFP allele (Fig. 1A and fig. S1, A and B). We tested the inducibility of the Pol II-GFP fusion protein by transient Cre expression in mESCs (fig. S1, C and D; see Supplementary Text) and the functionality by ChIP-seq (fig. S1, E to H). Next, we generated homozygous *Polr2b*^f/f^-GFP mice carrying hemizygous K14-Cre (*Polr2b*^f/f^-GFP; K14-Cre). Mice were viable and fertile with normal skin and hair follicles, and expressed keratinocyte-specific Pol II-GFP (Fig. 1, B and C, and fig. S2, A to D). Conventional ChIP-seq of *Polr2b*^f/f^-GFP; K14-Cre mouse back skin showed strong enrichment of known keratinocyte-specific genes in ChIP-seq with αGFP as compared to αPol II pull-down (fig. S2, E to I), as expected.

PReCIS-seq of mouse back skin from an adult skin resting stage (telogen) or a growth stage (anagen) with increased keratinocyte proliferation provided genome-wide nascent transcriptomic data (fig. S3, A to H’). Using previously published scRNA-seq data of whole skin at two developmental stages, we identified lineage-specific gene sets for: keratinocytes, mesenchymal, endothelial, and immune cells (fig. S4, A to C and data S1) (*25, 26*). Nascent transcript data in the αGFP pull-down revealed strong enrichment of engaged Pol II around the transcription start sites (TSSs) of keratinocyte lineage-specific genes (n=52), but not at genes specific to other lineages (Fig. 1D, fig. S4D, and data S2). In contrast, all gene sets displayed nascent transcript signal in the αPol II or input control conditions that probe all lineages in the skin (Fig. 1D, fig. S4D, and data S2). Specific examples for keratinocytes (i.e., *Krt14, Trp63*) and non-keratinocytes (i.e., *Cd74*) genes confirmed the expected signals (Fig. 1E) (*20*). Notably, the promoters of active genes showed divergent transcription, and transcription in the direction of the gene covered the entire transcription unit (Fig. 1E). Thus, PReCIS-seq provides high-resolution transcriptomic data of a specific cell lineage in its natural tissue microenvironment, without cumbersome and perturbing cell isolation procedures.

### Transcription regulation by Pol II promoter-recruitment vs. pause-release

Pol II engagement and nascent transcript studies in cultured and isolated cells documented transcription activation by two mechanisms: *de novo Pol II promoter-recruitment* or *Pol II promoter-proximal pause-release (3, 11, 13, 27)*. Specifically, upon transcriptional activation the ratio of normalized promoter-proximal counts (PPC) to gene body counts (GBC), known as the pausing index (PI), remains relatively constant on genes regulated mainly at promoter-recruitment and decreases on genes regulated at the pause-release step (Fig. 2A). Comparison of PReCIS-seq data of adult skin keratinocytes at telogen and anagen (a quiescence to proliferation transition) demonstrates distinct and reproducible differences and provides a genome-wide view of the transcription regulatory mechanisms operating during this cell state transition (Fig. 2A and fig. S5A). Cumulative genome-wide analyses show similar PI, PPC, and GBC distributions at the two stages analyzed (Fig. 2B, data S3, and Materials and Methods); however, by DESeq2 analysis (*28*), we identified 186 transcriptionally activated genes [GBC Fold Change (FC)>2], of which ∼21% (n=39) underwent increased Pol II pause-release (PI FC < -2) whereas ∼79% (n=147) displayed increased Pol II promoter-recruitment (PI ∼ constant) (Fig. 2, C and D). Function-based analysis showed an overall enrichment in cell cycle and signaling categories (Fig. 2E, fig. S5, B to E, and data S4), as expected at this quiescence-to-proliferation transition (*29*). Interestingly, HOMER analysis (*30*) in the 1 kb promoter region of these genes for transcription factor (TF) binding motifs showed that only promoter-recruitment– but not pause-release– regulated cell cycle genes contained motifs for TFs implicated in cell cycle gene regulation (*31-33*) (Fig. 2F and data S5). Several Shh signaling genes were upregulated at anagen (fig. S5E), as expected (*34*), and they also displayed both transcription activation mechanisms (fig. S5E). These data illuminate the global landscape of transcription activation mechanisms in a particular cell lineage within its intact tissue during adult mouse homeostasis.

**Fig. 2.**
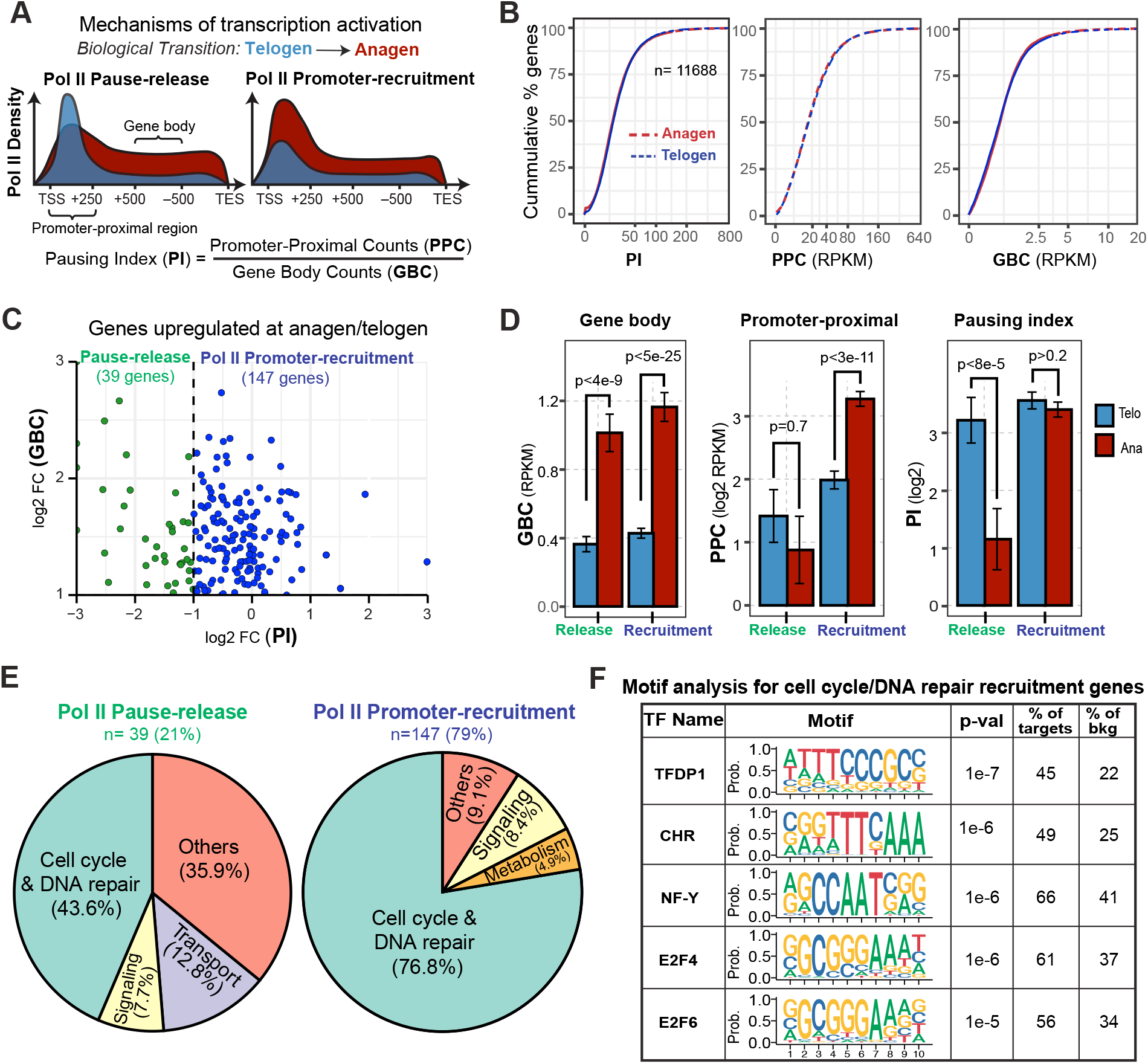
PReCIS-seq uncovers gene sets regulated by Pol II pause-release vs. promoter-recruitment. (**A**) Schematic of PReCIS-seq pattern expected for transcription activation via Pol II pause-release vs. promoter-recruitment during telogen to anagen transition. PI was calculated using RPKM normalized counts. (**B**) Cumulative distribution plots comparing PI, PPC and GBC in keratinocytes at telogen and anagen for all expressed genes. (**C**) Scatter plot showing changes in PI (x-axis) of transcriptionally upregulated genes (i.e., GBC FC > 2) (y-axis) at anagen. A cutoff of PI FC < -2 was used to delineate pause-release vs. promoter-recruitment genes. (**D**) Box plots comparing GBC, PPC and PI of the two gene categories regulated by pause-release (n=39) or promoter-recruitment (n=147). Mann-Whitney U rank sum test. **(E)** Pie charts summarizing distribution of top functional gene categories. **(F)** Summary of enriched transcription factor (TF) motifs from HOMER analysis on cell cycle and DNA repair genes regulated by promoter-recruitment.

### Low-paused lineage identity and high-paused cellular safeguarding genes

Development, cell cycle, and signaling genes can be “poised” for transcriptional activation, with high Pol II promoter-proximal pausing and pause-release regulation, as reported in Drosophila development and mammalian cell culture (*12, 35-37*). Little is known about genome-wide pausing of functional gene categories in a lineage of an intact mammalian tissue. PReCIS-seq of keratinocytes at telogen features the global status of Pol II engagement at a resting stage of skin homeostasis, which characterizes most of adult mouse life. We began the analysis by curating six gene sets expressed in keratinocytes with known biological functions (see Material and Methods): 1) keratinocyte-specific lineage identity genes (n=49); 2) development (n=34); 3) signaling (n=36); 4) metabolism (n=56); 5) cell cycle (n=105); and 6) cytoskeleton (n=58) (data S6). Interestingly, keratinocyte lineage identity genes were uniquely distinguished by high expression (indicated by high GBC) and very low pausing index (PI) (Fig. 3A and fig. S6A).

**Fig. 3.**
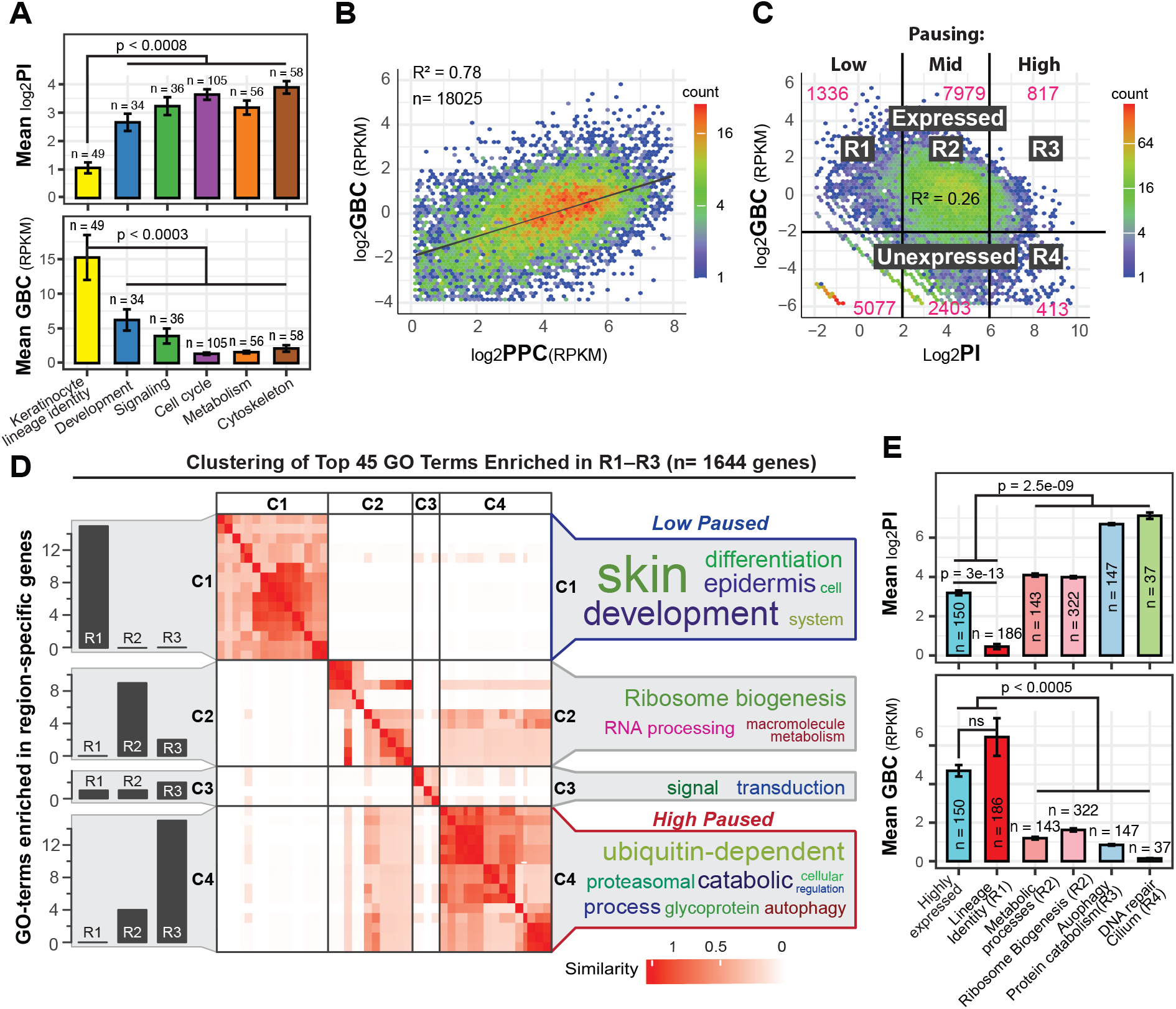
Genome-wide Pol II pausing distribution of telogen keratinocytes in intact skin tissue. **(A)** Bar plots of mean PI and mean GBC of six gene sets curated based on major biological functions (statistical analysis by ANOVA followed by Tukey HSD). **(B)** Comparison of gene body counts (GBC) vs. promoter-proximal counts (PPC) of all n=18,025 genes. **(C)** Comparison of gene expression (GBC) vs. pausing index (PI) and divisions of gene groups (see Materials and Methods) to obtain low- (R1), mid- (R2), and high- (R3) paused and expressed genes. R4, high-paused, unexpressed genes. **(D)** Binary cut clustering of top 45 unique GO terms from R1-R3 gene groups. Resulting C1-C4 clusters with bar plot distribution of R1-R3 genes (left panel), similarity matrix (middle panel), and most common GO words (right panel). **(E)** Bar plots of PI and GBC for the genes of top unique GO terms from R1-R4 gene groups and control of random highly expressed genes (statistical analysis by ANOVA followed by Tukey HSD).

Further unbiased assessment of genome-wide Pol II activity revealed positive correlation of gene expression (GBC) with Pol II promoter-proximal density (PPC), and a negative correlation with Pol II pausing index (PI) (Fig. 3, B and C). Genes expressed above the background (n=10,132) displayed various levels of pausing as follows: ∼79% mid-paused (n=7979 genes, R2 region in Fig. 3C), ∼13% (n=1336, R1) low-paused, and ∼8% (n=817, R3) high-paused (Fig. 3C and data S7; see Material and Methods). Finally, a small group of 413 genes were unexpressed (log2 GBC [RPKM] < -2) but highly paused (Log2 PI > 6) (Fig. 3C, R4). To functionally characterize differentially paused genes, we performed gene ontology (GO) analysis of expressed genes in the low-, mid-, and high-paused groups (R1-R3) (fig. S6B and data S8). Clustering of top unique GO-terms (total 45) using a binary cut method (*38*) demonstrates that low-paused genes are enriched in skin development and epidermis differentiation-associated keratinocyte lineage functions; mid-paused genes are enriched in ribosome biogenesis, RNA processing and macromolecule metabolism; and high-paused genes are enriched for ubiquitin-dependent protein catabolic processes and autophagy (Fig. 3D). Finally, high-paused unexpressed genes (R4) were enriched in DNA repair and cilium functions (fig. S6B). The average PIs of five gene categories derived from the top unique GO terms of groups R1-R4 confirmed these results and showed that high expression and low pausing are not necessarily correlated (Fig. 3E). Altogether, these analyses reveal the relationship of global Pol II pausing and gene expression status with gene sets of specific functions in an adult lineage of an intact mouse tissue (see Discussion).

### Enhancer proximity and promoter TF motifs relate to Pol II activity and pausing level

Pol II activity is influenced by transcription regulatory elements that contain TF binding sites and can act from a distance (e.g., enhancers) or close (e.g., promoters) to a gene TSS (*39*). While inspecting PReCIS-seq patterns of lineage-specific genes (i.e., *Krt14*), we noticed multiple divergently transcribed elements (Fig. 4A) reminiscent of active enhancers (*40*). To assess the capability of PReCIS-seq methodology to detect putative active enhancers genome-wide, we employed previously described dREG analysis (*41*) on our PReCIS-seq data at telogen. After exclusion of annotated promoters, we identified 5132 dREGs as putative enhancers, which showed divergent pattern of transcription from a TSS not associated with a NCBI RefSeq transcript (Fig. 4, B and C, data S9 and see Methods). These dREGs also showed high chromatin accessibility in published ATAC-seq data of epidermal keratinocytes at telogen (*42*) (fig. S7, A and B). These putative enhancers also displayed high overlap with mouse candidate cis-Regulatory Elements (cCREs) (*43*), of which a majority are classified as distal or proximal Enhancer Like Sites (dELS or pELS) (Fig. 4D and data S10). Importantly, analysis of enhancer-promoter proximity revealed a striking positive correlation with PReCIS-seq gene body transcription level, with most high expressed genes containing an active enhancer within ∼10kb of TSS whereas most medium and low expressed genes have active enhancers at ∼100kb or further away from TSS (Fig. 4E, fig. S7, C and D and data S11). We found only a mild (if any) inverse correlation with Pol II pausing level as defined by PI obtained from our PReCIS-seq telogen data (Fig. 4, F and G, fig. S7, E and F, and data S12 and S13).

**Fig. 4.**
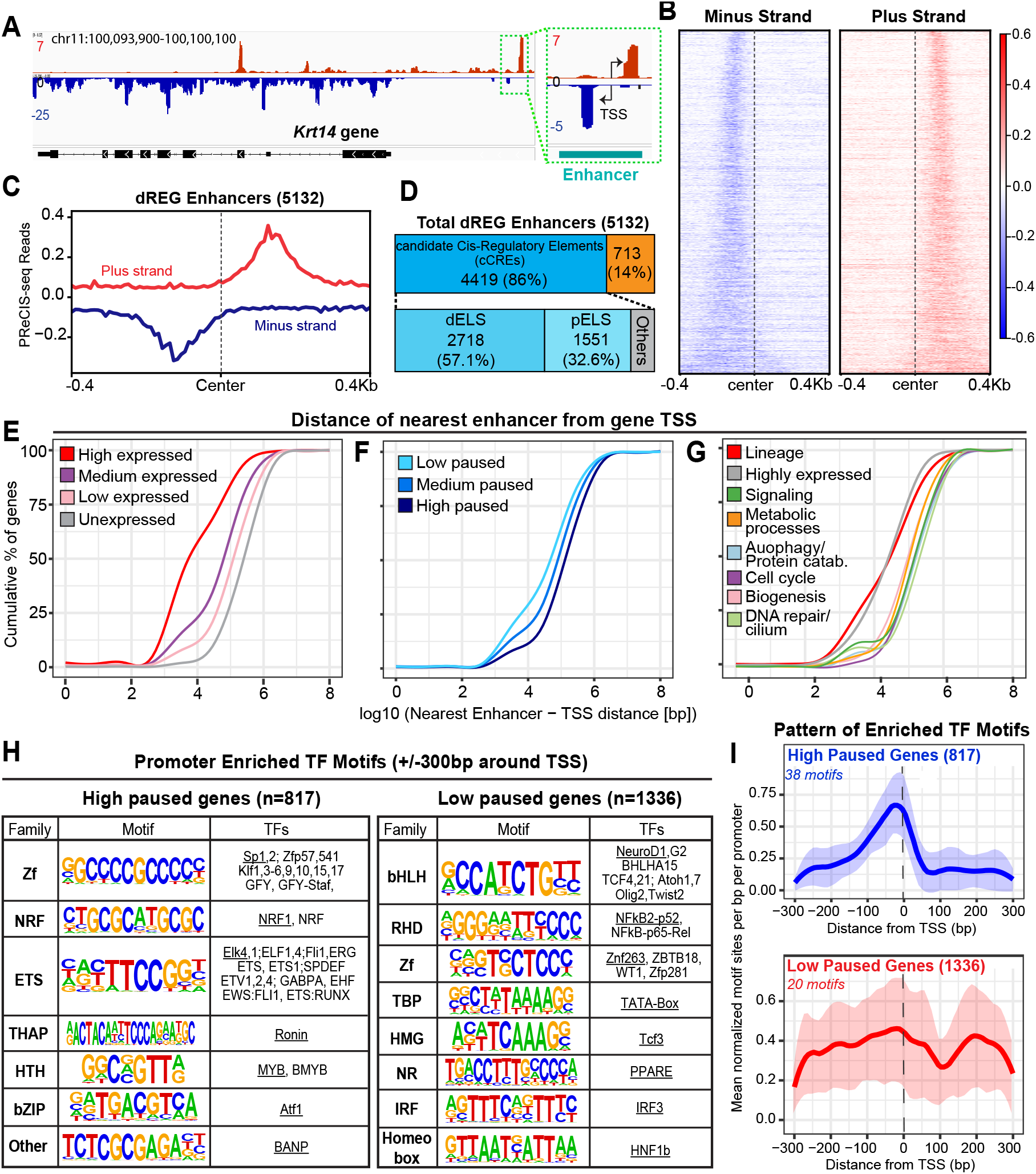
Active enhancer and promoter analysis reveals basic rules of Pol II activity. **(A)** Genome browser view of PReCIS-seq signal at the *Krt14* gene locus including an upstream putative enhancer displaying divergent transcription. **(B)** Heatmap of PReCIS-seq signal on minus (left panel) and plus strands (right panel) at dREG elements that are aligned at their center and sorted based on their size, shorter dREG elements on top and longer elements at the bottom. Dashed lines delineate the center of the divergent peaks. **(C)** Profile plot of plus and minus strand reads for all dREG elements (5132) showing their divergent transcription pattern, a typical characteristic of active enhancer elements. All elements were aligned at the center of divergent peaks. **(D)** Fractions of overlapped and non-overlapped dREG elements with candidate Cis Regulatory Elements (cCREs) from the ENCODE database. Most overlapped dREG elements are distal or proximal Enhancer Like Sites (dELS or pELS). **(E-G)** Cumulative distribution plot comparing the distance of the nearest enhancer for genes in different gene groups based on expression levels (E), PI levels (F) and GO functions (G). **(H)** Table summarizing differential enrichment of TF motifs in low and high paused gene promoters. TFs are grouped in TF families and a representative motif (underlined) for the top enriched TF is included in the table. **(I)** Metaplots depicts pattern of localization for TF motifs enriched in high and low paused genes relative to TSS for each gene group indicated. Shaded area represents mean ±SD of the normalized motif counts.

We then asked if specific TFs acting on either enhancer or promoter regions may be associated with Pol II promoter-proximal pausing level by comparing the low-paused and high-paused genes identified in our PReCIS-seq telogen data. TF DNA binding motif enrichment analysis using HOMER did not reveal significant enrichment within nearest enhancers of any gene group analyzed (Table S1). In contrast, multiple TF DNA binding motifs distinguished the promoter (+/-300bp around TSS) of low-vs. high-paused gene groups (Fig. 4H and data S14) and this was independent of expression level (Table S1). Some of the motifs associated with high pausing were also enriched in specific functional gene categories, such as Autophagy/Protein Catabolism when compared with the rest of the genome (Table S1). Interestingly, integration of localization patterns for all TF motifs identified in the high-paused gene group revealed strong positioning directly on the TSS or slightly upstream (Fig. 4I and fig. S8A). In contrast, TF motifs associated with low-paused genes showed on average a notable peak located more than 100bp downstream of TSS (Fig. 4I and fig. S8B, green arrows). In summary, these findings highlight the applicability of PReCIS-seq methodology to map active enhancers simultaneous with gene expression and Pol II pausing status; they revealed several basic rules of Pol II activity and pausing in a specific lineage of an intact tissue (see Discussion).

### Context-dependent dynamic Pol II pause-release of specific functional gene sets

Finally, we wondered if specific gene categories maintain their pausing patterns stably across different developmental and ‘*ex vivo*’ conditions. To this end, we examined the status of Pol II pausing for three intact-tissue, or ‘*in vivo’*, conditions [i.e., PD21, telogen; PD24, anagen; and embryonic day (E)16.5] and two ‘*ex vivo’* conditions [i.e., freshly isolated and cultured keratinocytes] (see Materials and Methods). Interestingly, nascent transcript data indicates comparable expression levels (GBC) but distinct proximal-promoter levels (PPC) and pausing statuses (PI) across conditions (fig. S9A, and data S3 and S15). To normalize across experimental conditions, we used several functional gene categories identified at telogen and plotted PI and GBC relative to the whole genome (Fig. 5, A to C’ and fig. S9, B to E’). Statistical pairwise analysis demonstrates context-dependent changes of pausing levels across conditions that were characteristic to each gene category (Fig. 5D and fig. S9F). Specifically, genes involved in tricarboxylic acid cycle and amino acid ‘metabolic process’ mid-paused at telogen (Fig. 3, R2) strongly maintained their pausing status across all experimental conditions (Fig. 5, A to A’ and D). In contrast, keratinocyte lineage identity genes low-paused at telogen (R1) gained pausing whereas high-paused autophagy and protein catabolism genes (R3) lost pausing in the *ex vivo* conditions (Fig. 5, B to D). Finally, biogenesis genes, cell cycle, and signaling genes mid-paused at telogen (R2) as well as high-paused, unexpressed DNA repair and cilium genes (R4) display minimal changes in pausing status across conditions (Fig. 5D and fig. S9, B to E’). Despite these changes, the low-paused (R1) gene group remained highly enriched in various keratinocyte lineage identity functions, irrespective of condition (Fig. 5E, fig. S9G and data S16). In contrast, the high-paused and expressed (R3) genes maintain top enrichment in autophagy and protein catabolism processes only in the intact tissue (Fig. 5E and fig. S9G). Instead, ‘signaling’ and ‘oxidative stress’ or ‘cell cycle’ and ‘DNA repair/cilium’ were top enriched in the high-paused and expressed genes *ex vivo*. Together, our data highlights the dynamic Pol II pausing status of functional gene groups within an intact mammalian-tissue environment and its profound change in *ex vivo* stress conditions (see Discussion).

**Fig. 5.**
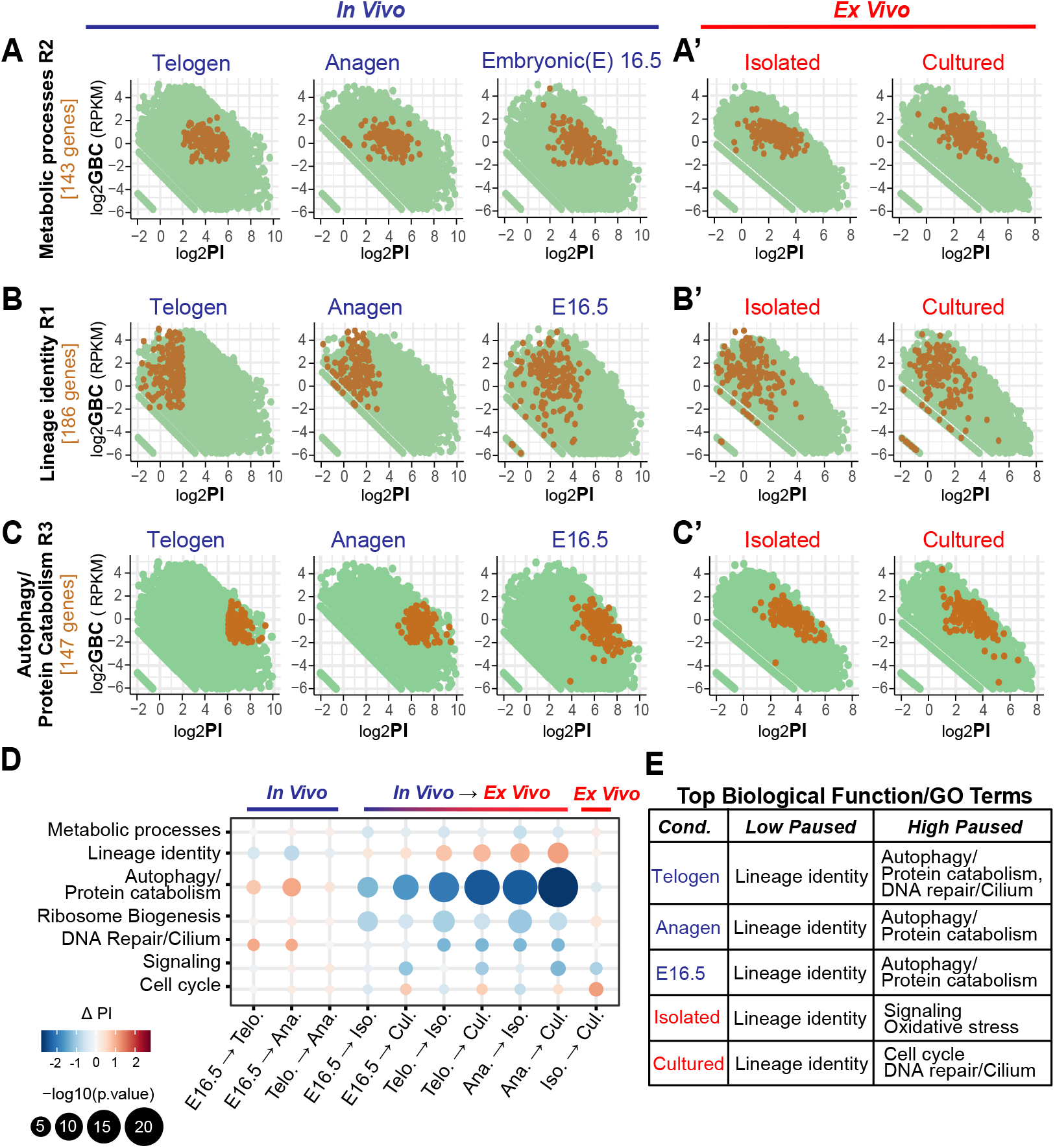
Context-dependent dynamic pause-release of specific gene categories. **(A-C’)** Scatter plots of GBC vs. PI for *in vivo* and *ex vivo* (isolated and cultured) experimental conditions. Specific gene categories indicated on y axis identified at telogen (see Fig. 3) are colored brown and the remaining genes in the genome are green. **(D)** Dot plot of the change in PI of specific gene categories as compared to the rest of the genome across pairwise comparisons for all five experimental conditions. **(E)** Table summarizing the top unique GO term(s) for low- and high-paused genes for all five experimental conditions.

### Considerations for future PReCIS-seq studies

Comparison of PReCIS-seq nascent transcript levels with steady-state mRNA levels measured by conventional RNA-seq could in theory provide information about the extent to which transcription versus mRNA stability contribute to gene expression at specific biological transitions. Using our *Polr2b*^f/f^-GFP; K14-Cre mice, we obtained total RNA-seq data of freshly isolated skin and GFP+ keratinocytes at telogen (fig. S10, A and B; see Methods). Direct comparison between PReCIS-seq gene body counts and RNA-seq counts shows moderate correlation independent of gene length (fig. S10, C and D). Interestingly, ∼5% of all detected genes show poor correlation between nascent transcription and total mRNA levels, suggesting regulation at the mRNA stability level for different functional gene categories for future exploration (fig. S10, E and F).

Although we focused here on adult skin keratinocytes, PReCIS-seq methodology can in principle be implemented to any mouse tissue where lineage-specific Cre recombinases are available. Since keratinocytes are a relatively abundant cell lineage, we tested the sensitivity limits of our method by employing the Shh-CreERT^2^ driver (*44*). This driver targets a small progenitor population in the hair follicle matrix, with labelled cells representing only ∼1% of the entire PD27 anagen skin tissue cells (fig. S11, A and B, and data S17). PReCIS-seq methodology and data analysis applied to skin of *Polr2b*^f/f^-GFP; Shh-CreERT^2^ mice demonstrate successful profiling of this small, targeted population (fig. S11, C to J and data S18). Although further optimizations and deeper sequencing of PReCIS-seq libraries are necessary for a more detailed and comprehensive analysis of this rare cell population, we are confident that our results demonstrate high sensitivity and broad applicability of PReCIS-seq for Pol II activity mapping in small cell populations within complex mammalian tissues.

## Discussion

Here, we developed PReCIS-seq technology to provide lineage-specific genome-wide quantitative mapping of transcriptionally-engaged RNA Pol II in a complex intact adult mouse tissue. These measurements provide views of Pol II as it progresses through critical regulatory steps of transcription initiation (Pol II promoter-recruitment) or elongation (Pol II pause-release); they also simultaneously map active divergent transcription sites at locations away from promoters that are likely to function as enhancers (*45, 46*). This approach can reveal fundamental mechanisms by which transcription factors and co-factors at promoters and enhancers regulate Pol II activity and gene expression during development, homeostasis and disease (*6, 47*). PReCIS-seq avoids laborious tissue dissociation and cell isolation procedures (*15, 16*), which can alter the spatiotemporal regulatory landscape. Using an adult keratinocyte prototype lineage in intact mouse skin at its resting stage (telogen), which characterizes most of mouse adult life (*29*), we uncovered several principles of gene regulation.

First, analysis of pausing index (PI) changes at specific biological transitions can reveal the critical step of transcription activation: Pol II promoter-recruitment vs. pause-release *(3, 11, 13, 27)*. Here, by focusing on a simple biological transition that involves increased proliferation in the keratinocyte lineage, we found several hundred activated genes implicated in signaling, cell cycle and DNA repair, as expected (*34*). Importantly, activated genes employed primarily Pol II promoter-recruitment with only about a fifth clearly utilizing Pol II pause-release mechanisms. Pol II pause-release regulation of cell cycle genes was also reported in an *in vitro* developmental context (*15*). Presumably, transcription factors and co-factors recruited to genes act on distinct steps in transcription regulation (*31-33*). Intriguingly, cell cycle related TF motifs decorated the promoters (within 1KB upstream of TSS) of Pol II promoter-recruitment but not pause-release genes, providing clear candidates for future mechanistic studies.

Second, Pol II pausing level on the proximal promoter inversely determine the rate of Pol II firing into elongation and the overall level of gene expression (*48*). High Pol II pausing is directly poising genes for rapid activation by pause-release (*12, 37, 49*). Previous functional studies that perturbed pause-release in various model systems, including the keratinocyte lineage of interest here (*50-52*), demonstrate the crucial biological relevance of this regulatory step (*11, 37*). Interestingly, we find that keratinocyte lineage-identity genes (i.e., related to keratinocyte differentiation, skin development, intermediate filament organization and wound healing (*21*)) show atypically low Pol II pausing in all intact-tissue and *ex vivo* conditions tested. These low-paused genes are also highly expressed, which demonstrates rapid firing from the Pol II paused site into effective elongation. On the other hand, the highest paused genes poised for activation in intact-skin keratinocytes were involved in cellular safeguarding processes, such as DNA break repair (*53*), autophagy - essential for recycling damaged organelles and maintaining healthy cells (*54*) - and the associated ubiquitin dependent protein catabolism (*55*). Indeed, these safeguarding processes are rapidly activated in adult skin keratinocytes upon acute UV exposure, an imminent stress constantly faced by this lineage (*56, 57*). Importantly, cellular safeguarding genes lose the highest-paused status upon tissue dissociation, likely due to increased Pol II firing into elongation in these new stress conditions (*48*). In fact, the highest-paused genes in *ex vivo* conditions were related to signaling, cell cycle, and oxidative stress, as reported in other cell culture studies (*37, 51, 58*). This likely reflects a high proliferative incentive from growth factors in the medium and new stress conditions outside the tissue hypoxic environment. These data pinpoint the dynamic context-dependent nature of pause-release and the importance of studying tissue regulatory mechanisms in the natural tissue context.

Third, multiple ON/OFF switch mechanisms influence Pol II pausing level, but how pause-release is differentially regulated on different promoters is unclear (*6-9*). Scarce literature reports attest the role of several developmental transcription factors (TF) in establishing pausing status or promoting pause-release on specific genes. These include the pioneering work on GAGA transcription factor for establishing paused Pol II (*59, 60*) and heat shock factor (HSF), TRIM28, MYC (bHLH), KLF4 and ZMYND8 (Zf), and MYB (HTH) transcription factors regulating pause-release (*61-64*). However, aside from these scattered examples, we currently do not have genome-wide consensus motifs or mechanistic rules connecting the rates of Pol II pause-release to specific transcription factors on promoters or enhancers. Interestingly, our analysis provides clear candidates of distinct TF consensus motifs associated with extreme levels of Pol II pausing. For example, bHLH and RHD factors are differentially enriched on low-paused gene promoters whereas NRF, ETS, and HTH factors preferentially decorate the high-paused gene promoters. Furthermore, we find that TF DNA binding motif positioning relative to the gene TSS differed greatly in the high-paused vs. low-paused gene groups. For the former, TFs bind directly on the TSS or immediately upstream. For the latter, specific TFs bind more than 100bp downstream of TSS beyond the typical Pol II pausing region, onto the previously reported inhibitory +1 nucleosome region (*65-67*). It is tempting to speculate that this binding might efficiently clear the way for rapid Pol II pause-release into elongation. Surprisingly, we found no enriched TF motifs associated with Pol II pausing on the nearest active enhancer elements. In addition, we found that relative distance of enhancers to promoters was associated with gene expression level but not with Pol II pausing. Future investigation is needed to understand how different classes of TFs act from specific positions on promoters and cooperate with enhancers to regulate the level of pause-release. In conclusion, we provide a new platform for precise mapping of Pol II activity and nascent transcription on genes and enhancers in specific cell lineages of intact mouse tissue to characterize regulation of biological transitions in their natural context. This work provides un-precedented access to basic mechanistic principles of gene regulation in development, homeostasis, and disease.

## Acknowledgments

We thank the Center for Animal Resources and Education (CARE) for mouse care and the Cornell University Biotechnology Resource Center (BRC) for their help with sequencing of various types of libraries; Christian Abratte and Rob Munroe from the Cornell University Transgenic Core Facility for their help with generation of the transgenic mice; Andrew Siefert from Cornell Statistical Consulting Unit for help with statistical analysis of gene dispersion presented in Fig. 5 and S9.

## Author contributions

GC and TT designed the experiments and wrote the manuscript. GC generated mice, developed and performed PReCIS-seq experiments and analysis, and all ChIP-seq and RNA-seq experiments. AW performed *ex vivo* PRO-seq. PV performed radioactive experiments. CB helped with embryo samples and statistical analyses. SH helped with experiments, figures, and writing methodology. SH and MD contributed to computational analysis. JR contributed to targeting vector design and ChIP. AO contributed to enhancer analysis. GC, JL and TT analyzed and interpreted all data. GC, SH, and TT prepared the figures. JL and SH edited the manuscript.

## Data and materials availability

All data are either available in manuscript or deposited in public databases. The ChIP-seq, RNA-seq and PReCIS-seq data have been deposited in the Gene Expression Omnibus (GEO) under accession codes GSE278914. All other data are available upon request from the corresponding author.

