## Supplemental information for "A lineage-specific nascent RNA assay unveils principles of gene regulation in tissue biology"

**The PDF file includes:**

Materials and Methods

Supplementary text

Figs. S1 to S11

References

**Other Supplementary Material for this manuscript includes the following:**

Data S1 to S19

Table S1

#### Materials and Methods:

##### Animals

All mouse work was performed in accordance with the Cornell University Institutional Animal Care and Use Committee (IACUC) guidelines (protocol no. 2007-0125). The K14-Cre mice were gifted from Elaine Fuchs' laboratory. The Shh-CreERT2 mice was obtained from Jackson laboratory (*44*).

##### Construction of a vector for targeting endogenous *Polr2b* locus in mouse ES cells.

To generate a conditional GFP-tagged RNA Polymerase II (*Polr2b*$^{f/f}$-GFP; K14-Cre) mouse line, we employed the CRISPR-Cas9 system to modify the 3'-end of the *Polr2b* gene, which encodes the second largest subunit of the RNA Polymerase II complex (RPB2). The targeting vector was designed as follows to replace the endogenous exon 25 with floxed exon 25, followed by the exon 25 fused GFP and 3x-FLAG. Precisely, a 4107 bp long targeting construct was designed that consists of a 635 bp long 5' homology arm followed by 1$^{st}$ LoxP site in the middle of intron 24, exon 25, a rabbit β-globin polyadenylation terminator signal (RBPA), 2$^{nd}$ LoxP site immediately after RBPA, partial intron 24, exon 25, linker sequence, 1$^{st}$ FRT site, linker, EGFP, 2$^{nd}$ FRT, linker, 3x FLAG with stop codon, and 1022 bp long 3' homology arm from 3' UTR of *Polr2b* gene. The targeting construct was partially synthesized by GenScript (www.genscript.com) and cloned in-house into pUC19. Next, guide RNA (gRNA) sequences targeting upstream and downstream of exon 25 were inserted into the pDG459 plasmid containing Cas9 (*68*). The sequence of gRNAs used are as follows: 5'-AGCGGGCAACGAATCATCTG-3' and 5'-GTTTAGCTGCCAGAAGTTCA-3'.

##### Electroporation of targeting construct and gRNA-Cas9 plasmid in mouse ES cells.

The targeting construct along with gRNA containing pDG459 plasmids were electroporated in V6.5 mouse ES cells by the Stem Cell and Transgenic Core Facility at Cornell University. The V6.5 mouse ES cells were grown to ~70% confluence on irradiated feeders in standard mouse ES cell medium containing LIF. Electroporation was performed using a Bio-Rad Gene Pulser II. 8x10^6 cells were electroporated in a 4mm cuvette with 25µg of targeting construct and pDG459 plasmid, at exponential program at 300V and 250uf resistance. Cells were briefly pipetted up and down after electroporation and replated onto 100mm gelatinized TC dishes. Initial selection was performed with the addition of 5 μg/ml puromycin to the culture medium for 48 hours, and then cells were cultured until individual clones could be manually picked (10 days). Picked clones were split into duplicate 96-well dishes for screening and cryopreservation.

**Screening of clones and validation of GFP tagged RNA Pol II expression in mouse ESCs**

The ES cells clones were first screened for the presence of GFP sequence in the genome using the 5'-AAGCTGACCCTGAAGTTCATCTGC-3' and 5'-CTTGTAGTTGCCGTCGTCCTTGAA-3' primers followed by using junction spanning primers as shown in fig. S1. The primers used for 5' junction screening were 5F 5'-TGGGAAACCTGCTCGCTGTAT-3', and 5R 5'-ATCTCAGTGGTATTTGTGAGCCAG-3', and for 3' junction were 3F 5'-ACTACAAAGACCATGACGGTGA-3', and 3R 5'-TTCCGCGGCAGCCAAAATTA-3'. Further, the entire integrated targeting construct insert in mouse ES cell genome was PCR amplified using different combinations of primers and verified by Sanger sequencing. To ensure Cre-inducible expression of RPB2-GFP fusion protein, selected clone (F8) of mouse ES cells were transfected with a modified Cre expressing plasmid under EF1a (EFS) promoter in pSECC vector (Addgene plasmid # 60820) (*69*), followed by selection of positive clones under neomycin and puromycin. The selected clones were further expanded and used for validation of GFP expression by confocal microscopy, FACS, and western blotting.

**Generation of RNA Pol II-GFP knock in mouse line and genotyping**

Positive clone F8 identified by screening was then recovered and expanded for ES cell microinjection. ES cells were microinjected into blastocyst stage embryos harvested from B6(Cg)-Tyrc-2J/J albino mice, and then transferred into pseudopregnant recipient females. ES cell incorporation into chimeras (F8 chimeras) was judged by coat color, and chimeras were bred to determine germline transmission. Offspring of F8 chimeras (knock-in line) were used to establish the *Polr2b*$^{f/f}$-GFP; K14-Cre mouse line. The following primers were used for genotyping of Pol II-GFP mice.

*Polr2b*$^{f/f}$-GFP -For 5'-GCCAGAAGTTCATGGTCAGATAC-3',
*Polr2b*$^{f/f}$-GFP; K14-Cre -For 5'-CATGGACGAGCTGTACAAGGC-3', and
*Polr2b*$^{f/f}$-GFP Common -Rev 5'-CGACAGCTATAACCCTGCTCTTG-3'.

The expected PCR product size for WT allele is 262 bp and for transgenic KI allele is 406 bp.

**Chromatin immunoprecipitation (ChIP)**

Dorsal skin tissue of *Polr2b*$^{f/f}$-GFP and *Polr2b*$^{f/f}$-GFP; K14-Cre mice were collected at PD24 and flash-frozen in liquid N2, followed by dry pulverization of frozen tissue through a cell crusher (https://cellcrusher.com/tissuepulverizer/). Samples were cross-linked with 1% formaldehyde, quenched with 125 mM glycine, and washed with ice cold PBS with protease inhibitors (ThermoFisher, A32965). After pelleting, samples were resuspended in ice-cold swelling buffer (25 mM HEPES pH 7.9, 1.5 mM MgCl2, 10 mM KCl, 0.1% NP40, 5 mM NaF, 2 mM Na2VO4, 1 mM PMSF, 1X PIC) and dounced with both tight and loose pestle for 20 times on ice, followed by filtering through a 70 µm filter. After centrifugation (1,000xg 10 min 4°C), samples were resuspended in sonication buffer (50 mM HEPES pH 7.9, 140 mM NaCl, 1mM EDTA, 1% Triton X-100, 0.1 % Na-Deoxycholate, 0.1% SDS, 5mM NaF, 2 mM Na2VO4, 1mM PMSF, 1X PIC) and fragmented using Bioruptor at high setting for 30 second ON + 30 second OFF pulses for a total of 30 minutes. The supernatant was collected after centrifugation (10,000xg 15 min 4°C). For pulldown, each sample was divided into three equal volumes followed by addition of 10 µL of Rabbit IgG (Cell Signaling, #2729), Mouse monoclonal anti-Pol II 8WG16 (Abcam, ab817), or Rabbit polyclonal anti-GFP (Abcam, ab290) antibodies and incubated overnight at 4°C with end-to-end rotation. Next, the samples were incubated with Dynabeads-Protein A (ThermoFisher 10002D) for 4 hours at 4°C. Samples were then washed with sonication buffer, high salt wash buffer (50 mM HEPES pH 7.9, 500 mM NaCl, 1 mM EDTA, 1% Triton X-100, 0.1 % Na-Deoxycholate, 0.1% SDS), LiCl wash buffer (20 mM Tris pH 8.0, 1 mM EDTA, 250 mM LiCl, 0.5% NP40, 0.5% Na-Deoxycholate), and TE buffer (10 mM Tris pH 8.0, 1 mM EDTA, 0.01% Tween-20). Samples were eluted by resuspending beads in elution buffer (50 mM Tris pH 7.5, 1 mM EDTA, 1% SDS) and thermomixed at 65°C for 30 minutes. Sample cross-linking was reversed by incubation with NaCl, EDTA and Proteinase K (160 mM, 5mM, 200 µg/mL respectively) overnight at 65°C. DNA was purified through phenol/chloroform extraction and precipitated using ethanol, sodium acetate, and glycoblue followed by centrifugation. The DNA pellets were then air dried and resuspended in TE buffer containing RNase cocktail (ThermoFisher, AM2286). DNA concentration was measured using the Qubit HS dsDNA kit (ThermoFisher, Q33238). DNA libraries were constructed by Tn5 tagmentation. Briefly, primer A 5′-TCGTCGGCAGCGTCAGATGTGTATAAGAGACAG-3’ or primer B 5′-GTCTCGTGGGCTCGGAGATGTGTATAAGAGACAG-3′ was annealed to Tn5-rev oligo 5′-

[phos]CTGTCTCTTATACACATCT-3′. Transposons were assembled using Tn5 and primers A and B in DF buffer (100 mM HEPES, pH 7.2, 200 mM NaCl, 0.2 mM EDTA, 2 mM DTT, 0.2% Triton X-100, 20% glycerol). Tagmented DNA was PCR amplified and purified with AMpure XP beads (Beckman Coulter, A63880) and quality checked by running on 6% Native PAGE gels and Fragment Analyzer. The ChIP DNA libraries were sequenced with Illumina NextSeq500, 2x37 bp, paired end, at BRC Genomics facility at Cornell University. Mouse ES cells were ChIP-seq profiled similarly, except chromatin were isolated from cultured ES cells instead of frozen skin tissue.

**Analysis of ChIP-seq data**

ChIP-seq reads were first trimmed with cutadapt (v1.18) (*70*), then mapped to the mouse reference genome mm10 with Bowtie2 (v2.3.5) (*71*). The BAM files were then converted to RPM normalized bigWig format and visualized using Integrated Genome Viewer (IGV) (*72*). The correlation analysis was performed using plotCorrelation from deepTools (v3.5.5) (*73*). Heatmap and profile plots of read signal across genes were generated using the plotHeatmap and plotProfile functions from deepTools. The epithelial lineage-specific genes were obtained from the HAIR-GEL (*74*) after filtering out non-keratinocyte genes based on their known expression pattern (cell-type specificity checked using protein atlas - https://www.proteinatlas.org).

**PReCIS-seq methodology:**

*Chromatin isolation from flash frozen tissue*

Back skin tissue was flash frozen immediately after harvest followed by dry pulverization using a cell crusher. Tissue samples were resuspended in ice cold swelling buffer with RNase inhibitors including SUPERase (ThermoFisher AM2696), Protector (Sigma-Aldrich, 3335402001) and RNasin™ Plus (Promega, N2615) RNase inhibitors. The samples were dounced twice both with tight and loose pestle, filtered (100-µm and 70 µm strainers) and centrifuged at 3000 rpm for 5 minutes at 4˚C. After removal of the supernatant, samples were incubated in 30 mL NUN buffer (20 mM HEPES-NaOH pH 7.5, 300 mM NaCl, 1 M Urea, 7.5 mM Mgcl2, 0.2 mM EDTA, 1% NP-40, 1 mM DTT, SUPERase, Protector, RNasin™ Plus) on ice for 30 minutes with intermittent vortexing (*75*). The samples were then centrifuged 4500 rpm for 10 minutes at 4˚C. The chromatin pellets were washed with ice-cold 50 mM Tris pH 7.5 (with RNase and protease inhibitors), and re-pelleted.

*Chromatin fragmentation and nascent RNA labelling by run-on reaction*

Following chromatin isolation, each sample was fragmented with DNase I (ThermoFisher, EN0523) at 37˚C with gentle mixing for 8 minutes. For run-on, samples were incubated with pre-warmed 2x run-on mix (10 mM Tris pH 8.0, 5 mM MgCl2, 300 mM KCl, 1 mM DTT, 1% sarkosyl, 40 µM biotin-CTP/UTP, 40 µM non-biotin-ATP/GTP with RNase and protease inhibitors) at 37 °C for 8 minutes with gentle mixing. The run-on reaction was diluted with a 10x dilution of ice-cold ChIP binding buffer (20 mM Tris pH 8.0, 150 mM NaCl, 1 % NP-40, 1 mM EDTA, 1x PIC, SUPERase In, Protector RNase Inhibitor, RNasin™ Plus) and centrifugation. The supernatant was pre-cleared using Dynabeads Protein A (ThermoFisher, 10002D), then incubated with 150µL of GFP coupled Protein A beads for 2 hours at 4˚C with rotation. After incubation, the beads were washed with ChIP binding buffer, Wash Buffer 1 (20 mM Tris pH 8.0, 150 mM NaCl, 1 % NP-40, 0.1 % SDS, 1 mM EDTA, 1x PIC, SUPERase In, Protector RNase Inhibitor), Wash Buffer 2 (20 mM Tris pH 8.0, 500 mM NaCl, 1 % NP-40, 0.1 % SDS, 1 mM EDTA, 1x PIC, SUPERase In, Protector RNase Inhibitor), and Wash Buffer 3 (20 mM Tris pH 8.0, 1 mM EDTA, 10% glycerol, SUPERase and Protector RNase Inhibitor).

*Isolation of nascent transcripts from the chromatin immunoprecipitated complexes*

Isolation of nascent transcripts and library preparation steps were followed as previously described for PRO-seq assay (*9*). Briefly, after thorough washing, nascent RNA was extracted from bead bound complexes using TRIzol (ThermoFisher, 15596026) and chloroform followed by centrifugation. TRIzol LS (ThermoFisher, 10296028) was used for input samples. Nascent RNA was precipitated with isopropanol and glycoblue. Input RNA was precipitated using 75% ethanol, resuspended in DEPC water, and passed through P30 gel columns (Bio-Rad, #7326250). Base hydrolysis, a standard step to fragment nascent RNAs in the conventional PRO-seq protocol, was omitted here as the size distribution of nascent RNAs was already in the appropriate range for generating libraries. Then, input RNA was conjugated to Streptavidin C1 beads (ThermoFisher 65001) and washed with high (50 mM Tris-Cl pH 7.5, 2 M NaCl, 0.5% Triton X-100) and low (5 mM Trix-Cl pH 7.5, 0.1% Triton X-100) salt wash buffers. TRIzol was added to the dried beads.

Starting this point onward, the treatments were applied to both input and nascent RNA samples. Chloroform was added to all samples, mixed, and samples were spun down. The top aqueous layer was collected, and RNA was precipitated with 100% ethanol and glycoblue as described above.

*3' Adapter ligation*

To ligate 3' adapters, RNA pellets were dissolved in 3' adapter solution followed by heat denaturation at 65˚C for 30 seconds and snap cooling on ice. 3' adapters were ligated to the nascent RNA using T4 RNA Ligase I (NEB, M0437M). To remove non-ligated 3' adapters, nascent RNA were captured on Streptavidin beads, followed by washing with the high and low salt wash buffers.

*On-bead 5' de-capping, hydroxyl repair and adapter ligation*

5' de-capping was carried out by first resuspending beads with nascent RNA in RppH reaction mix (10x ThermoPol buffer (NEB, B9004S), RppH (NEB, M0356S), SUPERase-In RNase Inhibitor) and incubating at 37°C for 45 minutes on thermomixer at 500 rpm. Following de-capping, 5' hydroxyl repair was conducted using T4 PNK (NEB, M0201L) at 37°C for one hour on thermomixer at 500 rpm. To ligate 5' RNA adapters, the beads were washed in C1 binding buffer and dissolved in the 5' adaptor solution. The beads were then heat denatured at 65˚C for 30 seconds and snap cooled on ice. 5' adapters were ligated to the nascent RNA using T4 RNA Ligase I (NEB, M0437M) in 10 µL reaction volume and 5 µM final 5'-adapter concentration. To extract nascent RNA, all samples were washed in high and low salt buffer before isolation with TRIzol and chloroform, and precipitation with ethanol and glycoblue.

*Reverse-transcription and amplification*

After centrifugation and air drying, RNA pellets were resuspended in a dNTP and reverse primer mix before denaturation at 65°C for 5 minutes and snap cooling. Then, reverse transcription was performed with Maxima RT enzyme (ThermoFisher, EP0752) with the following cycle settings: 15 minutes at 45˚C, 40 minutes at 50˚C, 10 minutes at 55˚C, 15 minutes at 70˚C. Samples were then test amplified and visualized on an 8% PAGE gel. Final amplification was performed through 13 cycles. Post-amplification DNA was purified with MinElute columns (Qiagen, 28004) and 1.5x AMpure XP beads. Library quality and size were checked using fragment analyzer and sequenced with NovoSeq6000 (NovoGene Inc, CA, USA) with 2x150bp paired-end settings. RP1 and RPI-n primers are from Illumina small RNA TruSeq design (Oligonucleotide sequences © 2015 Illumina, Inc. All rights reserved).

**PRO-seq profiling of isolated or cultured cells**

Newborn (PD 0.5) mouse skin keratinocytes were isolated as described previously (*76*) with slight modification to maintain cells at 4°C. Briefly, skin tissue was washed with 70% ethanol followed by PBS after decapitation. To separate the epidermis from the dermis, skin was incubated in cold Dispase at 4°C for overnight. The next day, the epidermis was separated from the dermis and incubated with 1% trypsin for 30 minutes at 4°C to prepare single cell suspensions. 20 to 30 million freshly isolated keratinocytes were either processed for PRO-seq library preparation as described previously (*77*) or cultured in vitro in low calcium (0.05 mM) containing E-media on irradiated mouse fibroblasts and then processed for PRO-seq library construction.

**Extraction of region-specific counts**

For PReCIS-seq libraries, adapters were trimmed using fastp (v0.23.2) (*78*). After trimming, reads were mapped to mouse rDNA using bowtie2 (v2.4.4) (*71*) to deplete nascent rRNA and to determine the fraction of total reads that map to rDNA. Then, the rRNA-depleted reads were mapped to the GRCm39 genome using STAR (v2.7.10b) (*79*) and UMI duplicate reads were removed by using umi_tools (v 1.1.2) (*80*). Deduplicated, mapped reads were processed from bam files to CPM-normalized bigwig files using samtools (*81*). The region-specific counts for promoter-proximal and gene body regions were obtained using BRGenomics R package (*82*).

**Classification of expressed and unexpressed genes of PReCIS-seq data**

To categorize genes into expressed and unexpressed categories based on PReCIS-seq signal in Fig. 3, the expression cut off was determined by i) background expression levels of 174 non-lineage (mesenchymal, endothelial and immune) genes (mean + SD) (background expression data for these genes are provided as supplementary data S7), ii) background expression levels on multiple un-annotated ~100 kb long non-gene regions, and iii) data distribution in scattered plot.

**Motif Analysis for upregulated cell cycle and DNA repair genes**

Motif enrichment analysis on promoters of cell cycle and DNA repair genes was performed using findMotifs.pl from HOMER software (*30*). Known motifs ranging from 8-12bp from 1kb upstream to 100bp downstream of transcription start sites were considered for enrichment analysis. Enriched motifs were sorted based on q-value significance, visualized using ggseqlogo (*83*) and ggplot2 packages in R.

**RNA-seq of FACS purified RNA Pol II-GFP$^+$ keratinocytes**

Dorsal skin of *Polr2b*$^{f/f}$-GFP; K14-Cre mice were collected at postnatal day (PD)21. A small piece of skin tissue was flash frozen (for RNA isolation from the whole skin tissue) and rest of the skin tissue was enzymatically digested using collagenase-trypsin method, as previously described (*84*). After preparing the single cell suspension, dead cells were stained with Propidium Iodide (PI). Live GFP$^+$ keratinocytes were FACS-purified and directly collected in TRIzol LS. Total RNAs were isolated following the manufacturer's protocol with the following additions: after the first phase separation, additional chloroform extraction step of the aqueous layer in Phase-lock Gel heavy tubes (Quanta Biosciences); addition of 1ul Glycoblue (ThermoFisher) immediately prior to isopropanol precipitation; two washes of the RNA pellet with 75% ethanol. Quality of each sample was confirmed by measuring RNA concentration using Qubit RNA HS kit (ThermoFisher) and by determining RNA integrity with Fragment Analyzer (Agilent). Ribosomal RNA was subtracted by hybridization from total RNA samples using the NEBNext rRNA Depletion Kit (New England Biolabs). TruSeq-barcoded RNAseq libraries were generated with the NEBNext Ultra II RNA Library Prep Kit (New England Biolabs) and libraries were sequenced on a NovaSeq with 2x150bp paired-end settings. Read preprocessing and alignment were performed using fastp (v0.23.2) (*78*) and STAR (v2.7.10b) (*79*). Adapters were trimmed using fastp (v0.23.2). Reads were mapped to the GRCm39 genome using STAR (v2.7.10b). Exon specific counts matrix was generated using featureCounts function from Rsubread (v2.14.2) followed by RPKM normalization for comparisons with nascent RNA-seq data.

**Immunofluorescence staining**

Immunofluorescence staining (IF) was performed following a standard protocol as described previously (*84*). Briefly, OCT-embedded samples were cryosectioned and fixed in 4%

paraformaldehyde for 10 minutes at room temperature (RT) followed by blocking in normal goat and donkey serum for 1 hour at RT. Blocking solution was then aspirated and samples were incubated with rabbit polyclonal Ki67 antibody (1:1000, Abcam, ab15580) and chicken polyclonal GFP antibody (1:100, Abcam, ab13970) for overnight at 4 °C. Next day, sections were washed with PBST and incubated with Alexa594-conjugated donkey anti-rabbit (Abcam, ab150076), or FITC-conjugated donkey anti-chicken secondary antibodies (Abcam, ab63507), and DAPI for one hour at RT. Then, the samples were washed and mounted with antifade and stored in -80°C before imaging. IF imaging was done using a fluorescent microscope (Leica DMI6000B) and digitally imaged using a fluorescence microscope camera (Leica K5) and the Leica Application Suite X software (v3.7.222383).

**Western blotting**

Proteins were isolated from cultured mouse ESCs or flash frozen mouse back skin tissue. Briefly, frozen tissues were dry pulverized using cell crusher (https://cellcrusher.com/tissuepulverizer/), followed by lysis in the ice-cold RIPA buffer (Sigma-Aldrich, R0278) containing protease inhibitors (Thermo Fisher Scientific A32965). Lysates were briefly sonicated and centrifuged at 16,000 RPM for 30 minutes at 4°C. Clear supernatant was collected and analyzed through SDS PAGE. Following antibodies were used to detect GFP tagged RPB2 and β-actin: (1:500, 50430-2-AP, Proteintech) and (1:10,000, MAB1501, Millipore), respectively.

**Single cell data analysis**

All datasets used in the single-cell RNA-seq analysis were obtained from NCBI Gene Expression Omnibus (GEO). Specifically for E16.5, scRNA-seq count matrix files (two replicates) were obtained from GEO GSE214441 (*25*), and for P21, scRNA-seq FASTQ files (three replicates) were obtained from GEO GSE153596 (*26*) and count matrices were generated using 10x Genomics Cell Ranger v.8.0.1 (*85*). Downstream data analysis was performed in R using the Seurat package version 5.1.0 (*86*). Cells that had between 500 and 5000 genes detected and had under 10% of the UMIs mapped to mitochondrial genes were retained for further analysis. After merging all samples, the transcript counts were log-normalized, and the data were integrated using *Harmony* (*87*). Genes enriched in each cell-type (cluster) were obtained by '*FindAllMarkers*' function using the Wilcox Rank Sum test.

**Lineage and functional gene sets curation**

To generate lineage-specific gene lists, in addition to well-known lineage markers like *Krt14* and *Krt10* for keratinocytes, and *Pecam1* and *Cdh5* for endothelial cells, we also obtained lineage-enriched genes from previously published single-cell RNA-seq datasets of whole skin (fig. S4, B and C). Genes enriched in keratinocytes, mesenchymal, endothelial and immune clusters were deemed "lineage genes" for the respective cell-type and used for enrichment analysis for PReCIS-seq data in Fig. 1. For functional gene categories, we obtained gene sets from publicly available resources such as KEGG (cell cycle - https://esbl.nhlbi.nih.gov/Signaling-Pathways/Cell-Cycle/, signaling - https://esbl.nhlbi.nih.gov/Signaling-Pathways/) as well as by literature searches.

**Gene ontology analysis and sorting of top unique GO terms**

Gene ontology analysis of different gene groups were performed using clusterProfiler package (v4.10.1) (*88*) with gene annotations org.Mm.eg.db (v3.18.0) database in R (v 4.3.3). The GO analysis results were sorted by adjusted p-value for further filtration of the significant GO-terms. Top 25 GO-terms from each group were used for clustering of similar GO-terms and their comprehensive representation in Fig. 3. The clustering and summarization were performed using binary cut method through simplifyEnrichment package (v1.12.0) (*89*) in R. To sort out top (significant) unique GO terms of low and high paused genes for all five conditions (Figs. 3 and 4), top unique GO terms were selected sequentially based on significance such that two unique GO terms have less than 70% overlap in genes.

**Enhancer analysis and motif search**

dREG elements were identified from telogen keratinocyte samples using mm39 mapped bigwig files (plus and minus strands) and dREG (*41*), which were then merged using BedTools (v2.26.0) (*90*). Mouse cCRE data (v4) was downloaded from ENCODE Portal (*43*). The mm10 coordinates of cCREs were converted to mm39 genome using liftOver program (*91*). Gene coordinates were derived from GENCODE mm39 GTF file, where all unique transcripts were selected. To filter candidate dREG enhancers, dREG elements that are over 500bp away from TSS of any GENCODE transcript are selected using BedTools intersect (v2.26.0). This resulted in 5132 candidate dREG enhancers (data S9).

Mouse keratinocyte ATAC-seq data were downloaded from GEO Database (GSM6840638) (*42*). These mm10 genome bigwig files were converted to mm39 genome using bigWigToBedGraph, liftOver, bedRemoveOverlap, and bedGraphToBigWig (v2.10) softwares. Heatmap and Metagene profiles of PReCIS-seq and ATAC-seq data were plotted using computeMatrix, plotProfile, and plotHeatmap functions from DeepTools (v3.5.2). In the heatmaps, dREG elements were aligned with their right boundaries (end with the higher genomic coordinate) and sorted based on element length. Data from 800 bp upstream and 200 bp downstream from the right boundary were plotted. Dashed lines represent the dREG element boundaries. All data in heatmap were plotted with the same dREG element order. Overlaps between dREG elements and cCREs were identified using BedTools intersect (v2.26.0). BedTools closest (v2.26.0) tool was used to determine the distance between dREG elements and gene TSSs.

Motif enrichment analysis on 300bp around gene TSS was performed using findMotifs.pl from Homer (*30*). The locations of individual enriched motifs were identified using annotatePeaks.pl in 10bp bins and plotted either individually or combined. Combined plots were generated to obtain pattern of motif localization around TSS. For that, value of total sites was scaled to have the maximum of 1 for each motif after combining both gene groups. Mean of scaled total sites per bin of all enriched motifs was calculated and plotted together.

**Statistics and reproducibility**

Statistical analyses were performed using R (v4.3.3). The RNA-seq and PReCIS-seq data were generated using two biological replicates for each method, one from a male mouse and one from a female mouse. To calculate the significance between two gene groups, unpaired two-tailed Student's t-test or ANOVA test were used, as appropriate, and multiple testing correction was done by the Benjamini method. For the analysis of genes groups distributions across conditions, the assessment of changes in distribution of specific gene group as compared to rest of the genes across four conditions (Embryo (E16.5), newborn, cultured, and adult keratinocytes) in multivariate framework of gene expression (gene body counts, Log2 of GB) and pausing index (Log2 of PI) was performed using the MANOVA.RM package, which doesn't assume normality or equal variance of variables between conditions. The MANOVA.wide function considers interactions between gene groups and conditions and tests whether the mean vectors of Log2 of GB and Log2 of PI differ significantly between conditions.

Pairwise comparisons of log2 PI means of different gene groups between two specific conditions were performed by calculating the contrast of group-adjusted means between the two conditions using the contrast function in the emmeans package (v1.10.3) in R. To calculate the difference in change of log2 PI of two different gene groups, a contrast of contrasts (generated from the previous step) was performed, also using emmeans. In the dot plots, contrast (or contrast of contrast) p-values were -log10 transformed and used for dot size scaling while contrast (or contrast of contrast) estimates were used to indicate the magnitude/direction of log2 PI change.

To calculate the significance of mean distances of nearest enhancers between two categories, a one-way ANOVA was first performed using the aov function in R to determine if there exists a significant difference of means between all categories. Post-hoc mean significance comparisons between two categories were then carried out using two-sample t-tests using the t-test function in R.

#### Supplementary text:

##### Functional validation of Pol II-GFP in mouse ES cells

Using CRISPR-Cas9 system, we targeted *Polr2b* allele that encodes for RPB2 - the second largest and one of the core subunits of RNA Pol II in mouse embryonic stem cells (mESCs). We designed a targeting vector carrying a floxed C-terminal exon (25) followed by a C-terminal exon fused with GFP and three copies of FLAG (fig. S1A, and Materials and Methods). Integration of the targeting allele at endogenous *Polr2b* locus was confirmed by 5' and 3' junction PCR (fig. S1B) and by sanger sequencing. Cre induced recombination of LoxP sites replaces an untagged exon 25 with GFP and 3x-FLAG tagged exon 25, further resulting in C-terminal tagging of RPB2 with GFP and 3x-FLAG. In this study, we will refer the unrecombined floxed allele as *Polr2b*$^{f/f}$-GFP*,* Cre induced recombined allele as *Polr2b*$^{f/f}$ -GFP; Cre, and the RPB2-GFP fusion protein as Pol II-GFP. To validate LoxP sites recombination induced Pol II-GFP expression, *Ef1a*-Cre (*69, 92*) was expressed in *Polr2b*$^{f/f}$ mESCs, and Cre dependent GFP expression was confirmed by western blotting and FACS analysis (fig. S1, C and D). Further, genome-wide validation of Pol II-GFP functionality was done by performing chromatin immunoprecipitation followed by sequencing (ChIP-seq) using antibody against C-terminal domain of RPB1 (αPol II) and GFP (αGFP) in *Polr2b*$^{f/f}$ and *Polr2b*$^{f/f}$-GFP; *Ef1a*-Cre mESCs (fig. S1, E to H). The ChIP-seq analysis showed almost identical genome-wide patterns for Pol II pulldown in *Polr2b*$^{f/f}$-GFP and *Polr2b*$^{f/f}$-GFP; *Ef1a*-Cre mESCs, suggesting that the Pol II-GFP functions similarly as WT Pol II. In addition, pulldown using GFP antibody in *Polr2b*$^{f/f}$-GFP; *Ef1a*-Cre cells showed even improved enrichment in gene body and towards the transcription end site (TES) as compared to Pol II antibody (known to preferentially enrich hypo-phosphorylated Pol II) and showed expected enrichment on target genes, highlighting unbiased enrichment of Pol II complexes from across the genomic regions using GFP pulldown (fig. S1, F to H).

##### Cre-dependent expression and functional validation of Pol II-GFP in skin keratinocytes

After generating the chimera mouse, we crossed *Polr2b*$^{f/f}$ mouse (further referred to as *Polr2b*$^{f/f}$-GFP) with keratinocytes specific Cre line (*Polr2b*$^{f/f}$-GFP; K14-Cre), in which only K14 positive cells (keratinocytes, ~20% from total skin cells) would express Pol II-GFP, whereas the rest of the cells in mouse will retain *Polr2b*$^{f/f}$-GFP allele. The Cre positive and negative littermates are indistinguishable (fig. S2A), suggesting that the GFP tagged version of RNA Pol II in K14 positive keratinocytes functions normally in vivo. Further, K14-Cre dependent and keratinocytes specific Pol II-GFP expression was confirmed at three different stages of mouse back development; embryonic day 16.5, morphogenesis at postnatal day (PD) 13 and quiescence phase at telogen (PD21), which showed that the expression of the tagged version is strictly dependent on the presence of Cre (fig. S2, C and D). Next to test cell-type specific chromatin enrichment potential of our methodology, we performed chromatin immunoprecipitation using Pol II and GFP antibodies on skin tissue from *Polr2b*$^{f/f}$-GFP and *Polr2b*$^{f/f}$-GFP; K14-Cre mice (fig. S2E). Our ChIP-seq data showed strong enrichment for keratinocytes specific genes over non-keratinocytes genes in GFP pulldown condition (fig. S2, F to I), confirming robust potential of our strategy for cell-type specific chromatin enrichment from heterogenous whole skin tissue and normal functionality of Pol II-GFP in vivo.

### Supplementary figures:

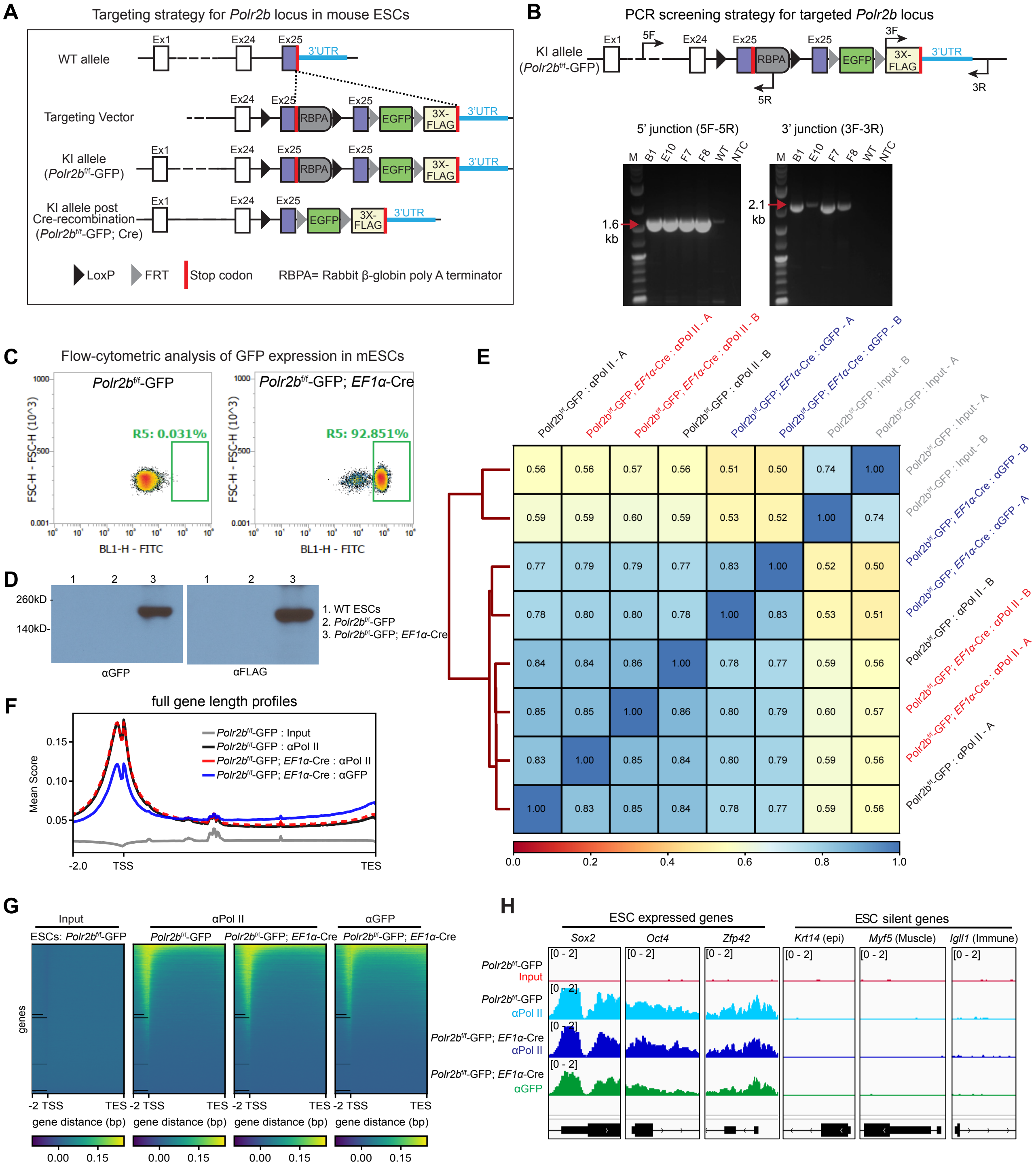

**Fig. S1. – Related to Fig. 1 – Characterization of Pol II-GFP expression in mouse ESCs and functional validation by ChIP-seq**

**(A)** Targeting strategy for *Polr2b* locus in mouse embryonic stem cells (mESCs) using CRISPR/Cas9. For a detailed description, see Materials and Methods. **(B)** PCR screening strategy for targeted *Polr2b* locus and validation of targeted insertion into mESC genome. **(C)** FACS of mESCs carrying the knock in (KI) allele demonstrates GFP expression in *Ef1α*-Cre-targeted mESCs. **(D)** Western blot of GFP and FLAG verifying expression of tagged-RPB2 in mESCs after *Ef1α*-Cre transfection (RPB2-GFP expected molecular weight is ~170 kDa). **(E)** Correlation matrix of ChIP-seq data of Pol II (control) and GFP pull-downs from *Polr2b*$^{f/f}$-GFP and *Polr2b*$^{f/f}$-GFP; *Ef1α*-Cre mESCs. **(F)** Profile plot of ChIP-seq data showing nearly identical pattern of Pol II pull-down across the gene length using Pol II control antibody for both *Polr2b*$^{f/f}$-GFP and *Polr2b*$^{f/f}$-GFP; *Ef1α*-Cre samples, indicating normal functionality of tagged Pol II. Note that GFP pull-down efficiently captures Pol II across the gene unit as compared to the Pol II antibody (8WG16), which preferentially captures hypo-phosphorylated Pol II that are enriched around the transcription start site (TSS). **(G)** Heatmaps of ChIP-seq data highlighting the observed enrichment pattern in (F) genome-wide. **(H)** Genome browser views of representative ESC expressed and silent genes in ChIP-seq data of *Polr2b*$^{f/f}$-GFP and *Polr2b*$^{f/f}$-GFP; *Ef1α*-Cre samples. Note strong enrichment of ESC genes and depletion of non-ESC genes in Pol II and GFP pull-down, as expected.

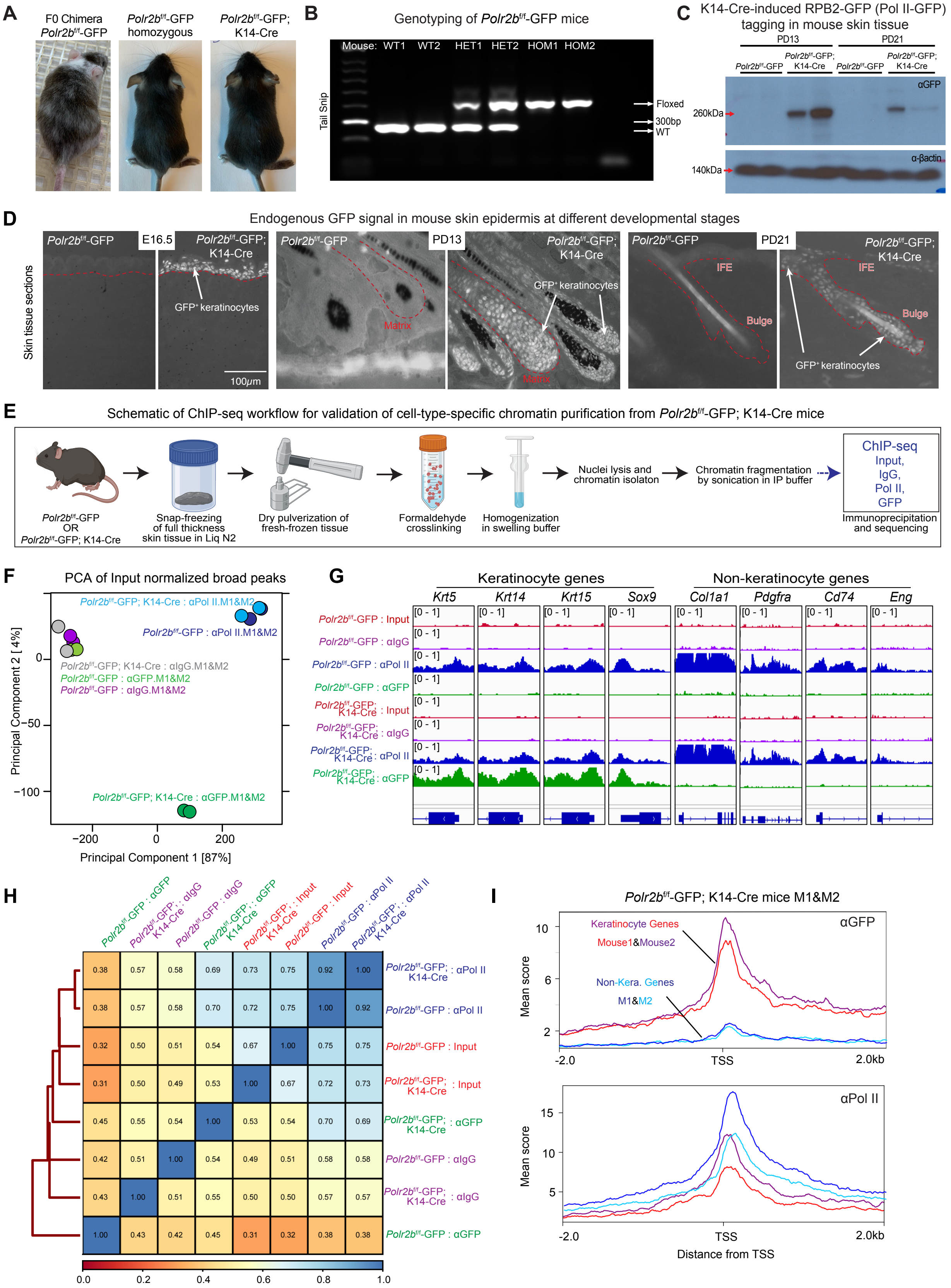
A
F0 Chimera Polr2b^f/f-GFP
Polr2b^f/f-GFP homozygous
Polr2b^f/f-GFP; K14-Cre
B
Genotyping of Polr2b^f/f-GFP mice
Mouse: WT1 WT2 HET1 HET2 HOM1 HOM2
Tail Snip
Floxed
300bp
WT
C
K14-Cre-induced RPB2-GFP (Pol II-GFP) tagging in mouse skin tissue
PD13
PD21
Polr2b^f/f-GFP
Polr2b^f/f-GFP; K14-Cre
αGFP
260kDa
140kDa
α-βactin
D
Endogenous GFP signal in mouse skin epidermis at different developmental stages
Skin tissue sections
E16.5
PD13
PD21
GFP⁺ keratinocytes
Matrix
IFE
Bulge
100µm
E
Schematic of ChIP-seq workflow for validation of cell-type-specific chromatin purification from Polr2b^f/f-GFP; K14-Cre mice
Polr2b^f/f-GFP OR Polr2b^f/f-GFP; K14-Cre
Snap-freezing of full thickness skin tissue in Liq N2
Dry pulverization of fresh-frozen tissue
Formaldehyde crosslinking
Homogenization in swelling buffer
Nuclei lysis and chromatin isolaton
Chromatin fragmentation by sonication in IP buffer
ChIP-seq Input, IgG, Pol II, GFP
Immunoprecipitation and sequencing
F
PCA of Input normalized broad peaks
Polr2b^f/f-GFP; K14-Cre : αPol II.M1&M2
Polr2b^f/f-GFP : αPol II.M1&M2
Polr2b^f/f-GFP; K14-Cre : αIgG.M1&M2
Polr2b^f/f-GFP : αGFP.M1&M2
Polr2b^f/f-GFP : αIgG.M1&M2
Polr2b^f/f-GFP; K14-Cre : αGFP.M1&M2
Principal Component 2 [4%]
Principal Component 1 [87%]
G
Keratinocyte genes
Non-keratinocyte genes
Krt5 Krt14 Krt15 Sox9 Col1a1 Pdgfra Cd74 Eng
Polr2b^f/f-GFP : Input
Polr2b^f/f-GFP : αIgG
Polr2b^f/f-GFP : αPol II
Polr2b^f/f-GFP : αGFP
Polr2b^f/f-GFP; K14-Cre : Input
Polr2b^f/f-GFP; K14-Cre : αIgG
Polr2b^f/f-GFP; K14-Cre : αPol II
Polr2b^f/f-GFP; K14-Cre : αGFP
[0 - 1]
H
I
Polr2b^f/f-GFP; K14-Cre mice M1&M2
αGFP
Keratinocyte Genes
Mouse1&Mouse2
Non-Kera. Genes
M1&M2
Mean score
αPol II
TSS
Distance from TSS

**Fig. S2. – Related to Fig. 1 – Functional characterization of Pol II-GFP homozygous mice through ChIP-seq and validation of cell-type specific chromatin enrichment *in vivo***

**(A)** F0 chimera bred to homozygosity, and normal phenotypic appearance of both *Polr2b*$^{f/f}$-GFP and *Polr2b*$^{f/f}$-GFP with hemizygous K14-Cre (*Polr2b*$^{f/f}$-GFP; K14-Cre) mice. **(B)** PCR genotyping for screening heterozygous and homozygous *Polr2b*$^{f/f}$-GFP mice. **(C)** Western blotting showing K14-Cre-dependent GFP expression in mouse skin at different developmental stages (PD - postnatal day). **(D)** Endogenous GFP fluorescence images of skin tissue sections showing nuclear, keratinocyte-specific GFP in the epidermis of *Polr2b*$^{f/f}$-GFP; K14-Cre mice at all developmental stages analyzed (IFE - interfollicular epidermis; scale bar, 100 μm; K14$^{+}$ epithelium is delimited with dotted red lines). **(E)** Schematic describing ChIP-seq workflow for validation of cell-type specific chromatin purification from *Polr2b*$^{f/f}$-GFP; K14-Cre mice. **(F)** PCA of input-normalized broad peaks of different ChIP-seq samples, showing clustering of Pol II pull-down samples from K14-Cre$^{+}$ and control mice, and clear separation of GFP pull-down samples from *Polr2b*$^{f/f}$-GFP; K14-Cre mice. M – mouse. **(G)** Genome browser views of ChIP-seq data for select keratinocyte and non-keratinocyte genes. In GFP pull-down in *Polr2b*$^{f/f}$-GFP; K14-Cre mice, keratinocyte genes were strongly enriched whereas non-keratinocyte genes were depleted. **(H)** Correlation matrix of ChIP-seq data of Pol II and GFP pull-down of *Polr2b*$^{f/f}$-GFP and *Polr2b*$^{f/f}$-GFP; K14-Cre mice. Pol II pull-downs from both genotypes showed high correlation, confirming normal functionality of Pol II-GFP *in vivo*. **(I)** Profile plots of ChIP-seq signal on 75 keratinocyte and 75 non-keratinocyte top expressed genes from scRNA-seq data of skin cells (See Materials and Methods) for two mice (M1 and M2). Note enrichment of Pol II-GFP signal on the TSS of keratinocyte genes and depletion on non-keratinocyte genes only in GFP pull-down.

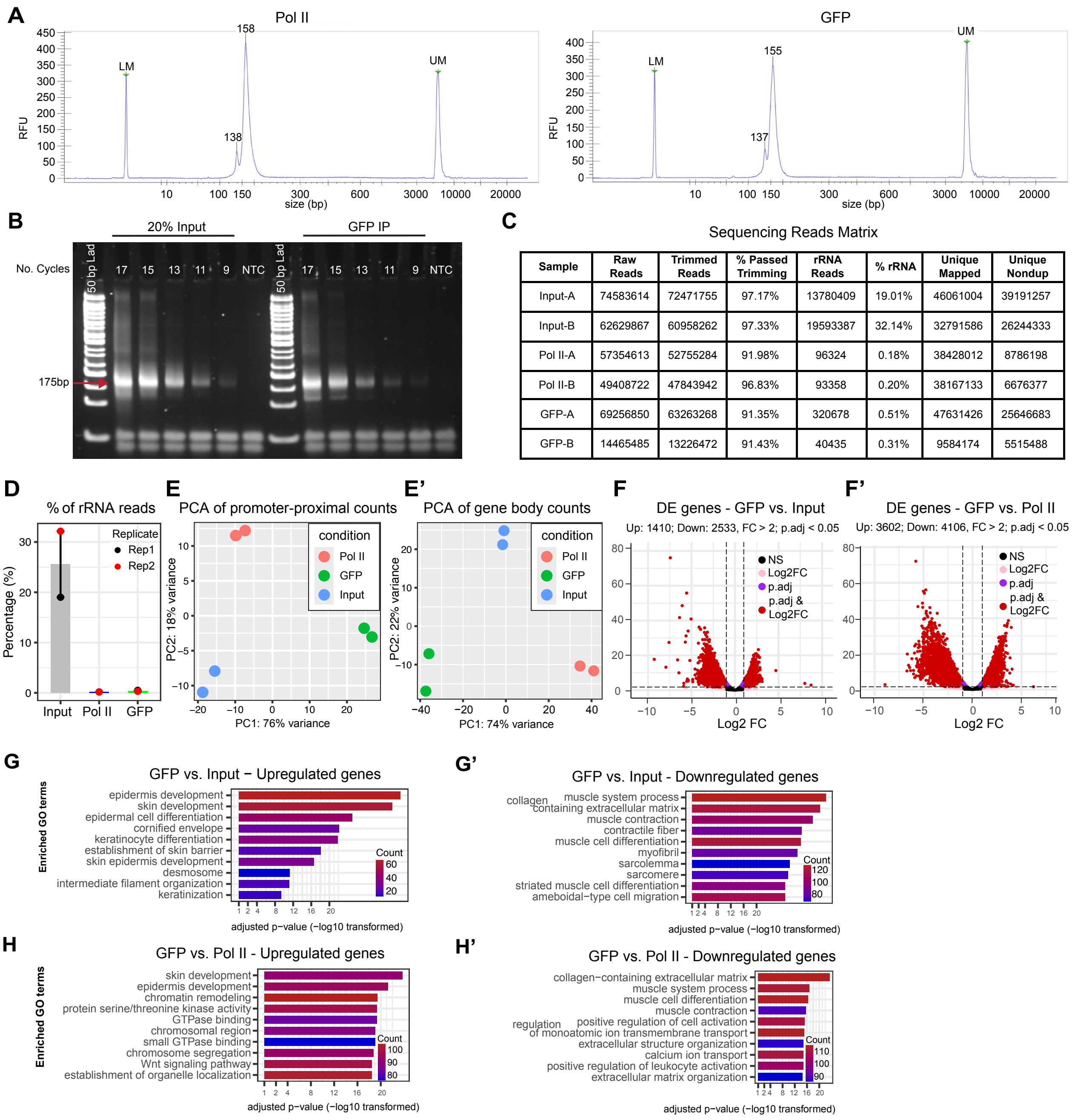


| Sample | Raw Reads | Trimmed Reads | % Passed Trimming | rRNA Reads | % rRNA | Unique Mapped | Unique Nondup |
|---|---|---|---|---|---|---|---|
| Input-A | 74583614 | 72471755 | 97.17% | 13780409 | 19.01% | 46061004 | 39191257 |
| Input-B | 62629867 | 60958262 | 97.33% | 19593387 | 32.14% | 32791586 | 26244333 |
| Pol II-A | 57354613 | 52755284 | 91.98% | 96324 | 0.18% | 38428012 | 8786198 |
| Pol II-B | 49408722 | 47843942 | 96.83% | 93358 | 0.20% | 38167133 | 6676377 |
| GFP-A | 69256850 | 63263268 | 91.35% | 320678 | 0.51% | 47631426 | 25646683 |
| GFP-B | 14465485 | 13226472 | 91.43% | 40435 | 0.31% | 9584174 | 5515488 |

**Fig. S3. – Related to Fig. 1 – Development and implementation of PReCIS-seq on mouse skin tissue at telogen**

**(A)** Bioanalyzer profile of PReCIS-seq libraries of Pol II and GFP pull-down samples (RFU - Relative Fluorescence Units). **(B)** PCR of PReCIS-seq libraries showing quality and fragment distribution. **(C)** Table summarizing mapped read counts before and after de-duplication. **(D)** Bar plot shows strong depletion of ribosomal RNA (rRNA) reads in Pol II and GFP pull-downs relative to input. **(E-E')** PCA plot showing high correlation of biological replicates (two mice) for promoter-proximal counts and gene body counts at telogen. **(F-F')** Volcano plots showing differentially expressed genes between GFP pull-down vs. input (F) and GFP pull-down vs. Pol II pull-down (F'). **(G-H')** Bar plots showing enriched Gene Ontology terms of significantly up- and down-regulated genes from F-F'. Upregulated genes in GFP pull-down show enrichment in skin and epithelium-specific processes as compared to input and Pol II pull-down.

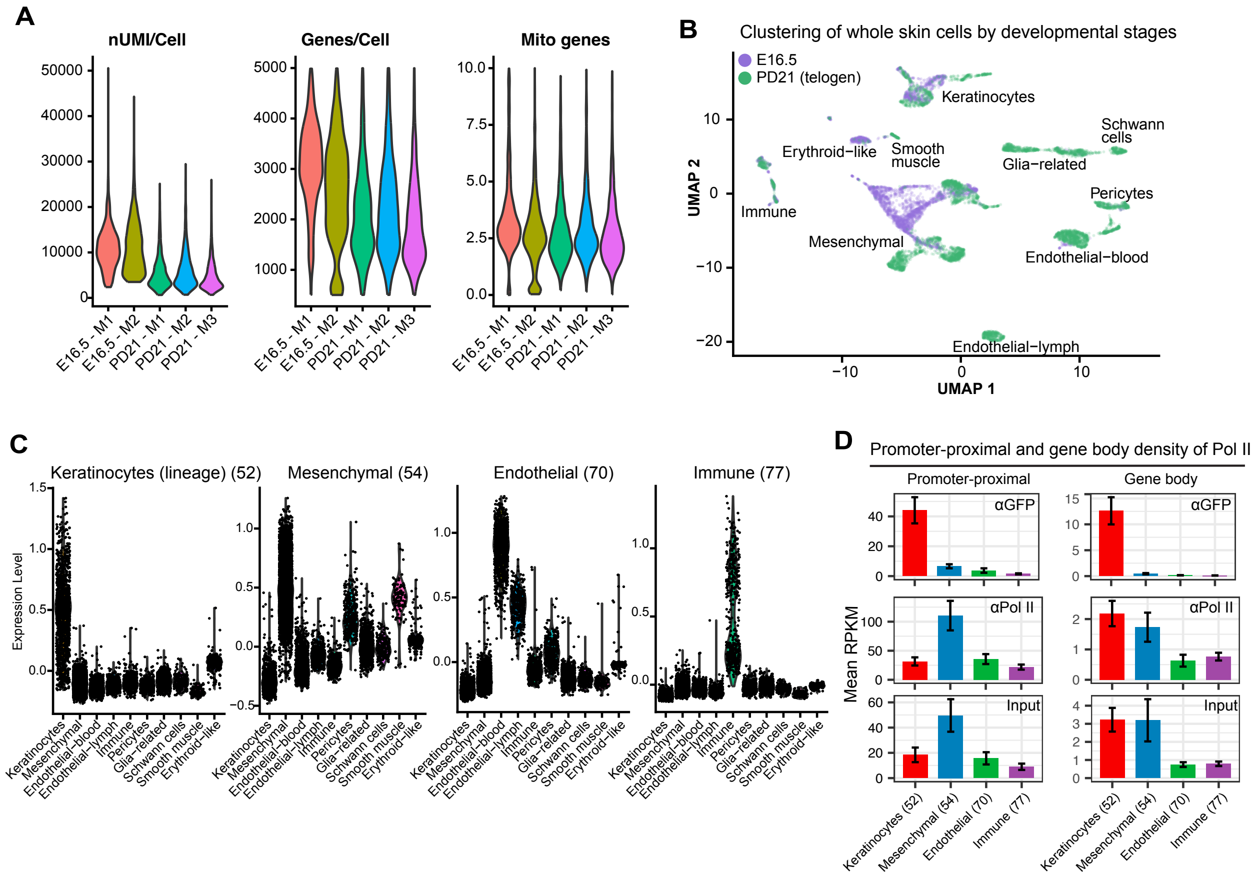


**Fig. S4. – Related to Fig. 1 – Cell-type-specific enrichment analysis of PReCIS-seq data**

**(A)** Quality of selected cells from two mouse replicates of E16.5 (M1, M2) and three replicates of PD21 (M1, M2, M3) subjected to scRNA-seq analysis extracted from (*25, 26*). nUMI/Cell – Unique Molecular Identifiers detected per cell, Genes/Cell – number of genes detected per cell, Mito genes – percentage of mitochondrial genes per cell, M – mouse. **(B)** Uniform Manifold Approximation and Projection (UMAP) plot of E16.5 and PD21 whole skin single-cell RNA-seq datasets. Samples from both stages were integrated using Harmony before plotting. Keratinocytes from both stages remain tightly clustered and distinct from all other non-epithelial cell types. **(C)** Violin plots showing enriched expression of lineage-specific genes in their respective single cell population. **(D)** Bar plots of mean RPKM-normalized reads of PReCIS-seq data of four skin lineage specific gene sets. Note more than 20-fold enrichment of keratinocyte-specific genes in GFP pull-down in both promoter-proximal and gene body regions.

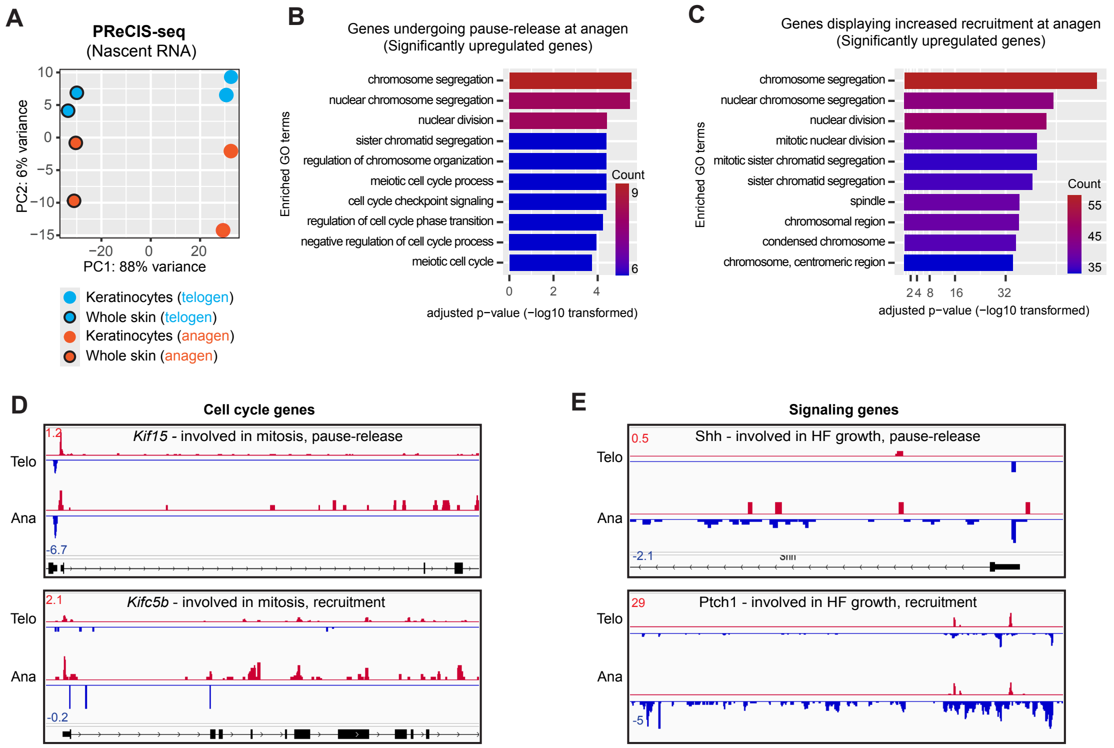


**Fig. S5. – Related to Fig. 2 – Classification of genes undergoing Pol II pause-release or displaying increased promoter-recruitment at telogen-anagen transition**

**(A)** PCA analysis of DESeq2 counts of PReCIS-seq data from GFP$^{+}$ keratinocytes and whole skin tissue at telogen and anagen. **(B)** Enriched GO terms of upregulated genes undergoing pause-release (PI FC < -2) at anagen. **(C)** Enriched GO terms of upregulated genes displaying increased promoter-recruitment without change in Pol II pausing at anagen. **(D)** Genome browser views of PReCIS-seq for cell cycle genes *Kif15* and *Kifc5b* displaying pause-release and promoter-recruitment regulation at anagen, respectively. **(E)** Genome browser views of PReCIS-seq for signaling genes *Shh* and *Ptch1* displaying pause-release and promoter-recruitment regulation at anagen, respectively.

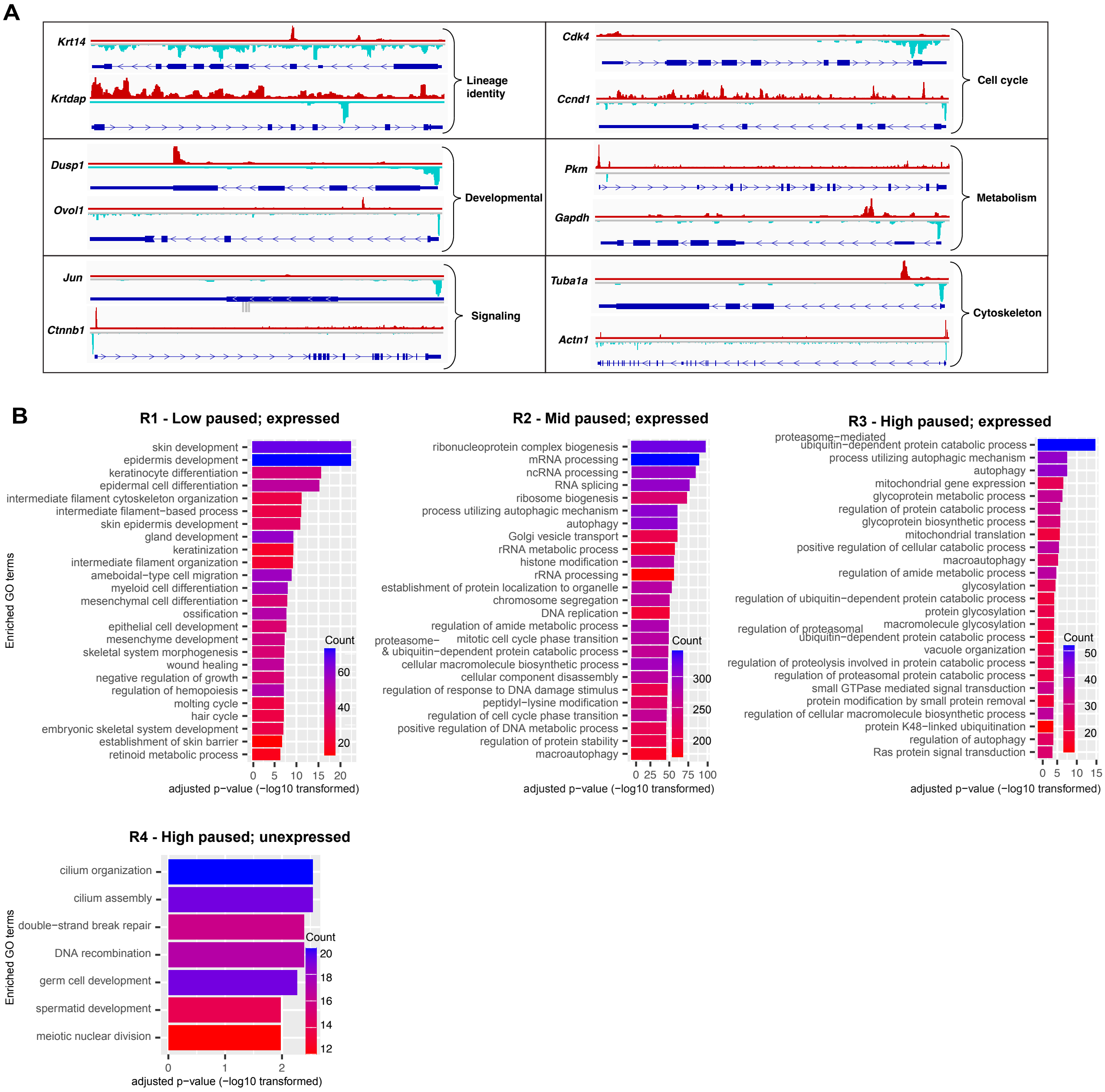


**Fig. S6. – Related to Fig. 3 – Pol II displays differential promoter-proximal pausing on different gene groups *in vivo***

**(A)** Genome browser views of telogen PReCIS-seq data showing occupancy of transcriptionally-engaged Pol II on select genes representing six major biological functions. **(B)** Bar plots of top 25 significantly enriched GO terms in each of the 4 gene groups as identified in Fig. 3C (See Data S8 for complete GO analysis results for each region).

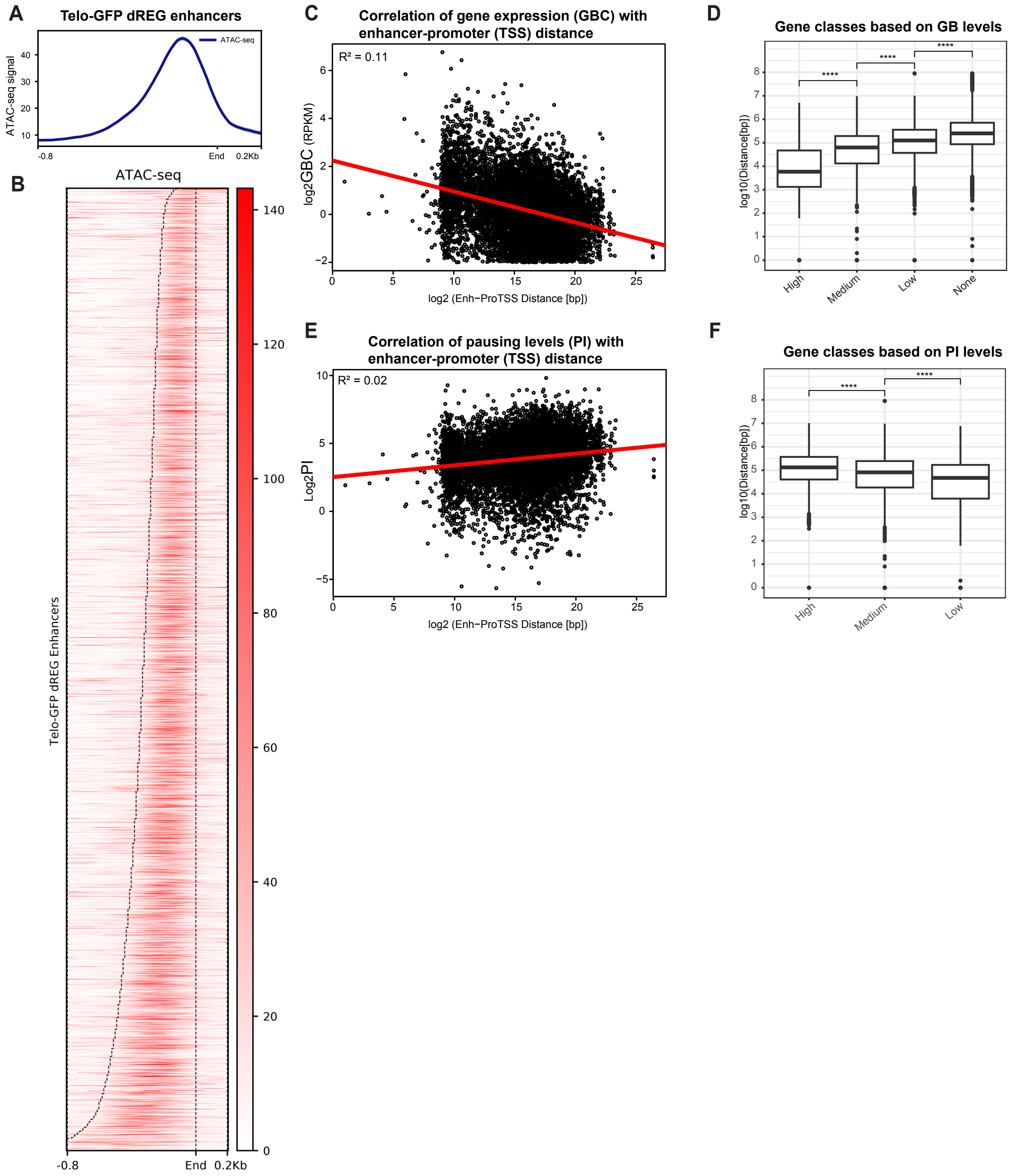
A
Telo-GFP dREG enhancers
ATAC-seq
ATAC-seq signal
40
30
20
10
-0.8
End
0.2Kb
B
ATAC-seq
Telo-GFP dREG Enhancers
140
120
100
80
60
40
20
0
-0.8
End
0.2Kb
C
Correlation of gene expression (GBC) with enhancer-promoter (TSS) distance
R² = 0.11
log2GBC (RPKM)
log2 (Enh−ProTSS Distance [bp])
D
Gene classes based on GB levels
log10(Distance[bp])
High
Medium
Low
None
E
Correlation of pausing levels (PI) with enhancer-promoter (TSS) distance
R² = 0.02
Log2PI
log2 (Enh−ProTSS Distance [bp])
F
Gene classes based on PI levels
log10(Distance[bp])
High
Medium
Low

**Fig. S7. – Related to Fig. 4 – Identification of putative enhancers and their association with Pol II activity**

**(A-B)** Profile plot and heatmap of ATAC-seq data showing chromatin accessibility at dREG elements (5132). ATAC-seq data was obtained from a previously published dataset in telogen keratinocytes (*42*). All dREG elements were aligned to the end position of the plus strand peak. Dashed line shows the left boundary of each dREG element. **(C)** Scattered plot showing correlation between level of gene expression (GBC) with enhancer proximity to gene TSS. Only expressed genes were considered for this correlation analysis. **(D)** Box plot showing mean distance of nearest enhancer for different categories of genes based on gene expression levels: High (Log2 GB (RPKM) >2), Medium (Log2 (RPKM) GB <2 & >0), Low (Log2 (RPKM) GB <0 & >-2), and unexpressed (Log2 (RPKM) GB <-2) genes. Two-sample t-test was used to calculate significance. **(E-F)** Scattered plot and Box plot same as in panel (C) and (D) for different categories of genes based on pausing index (PI): High (Log2 PI >6), Medium (Log2 PI >2 & <6), and Low PI (Log2 PI <2) genes. Two-sample t-test was used to calculate significance.

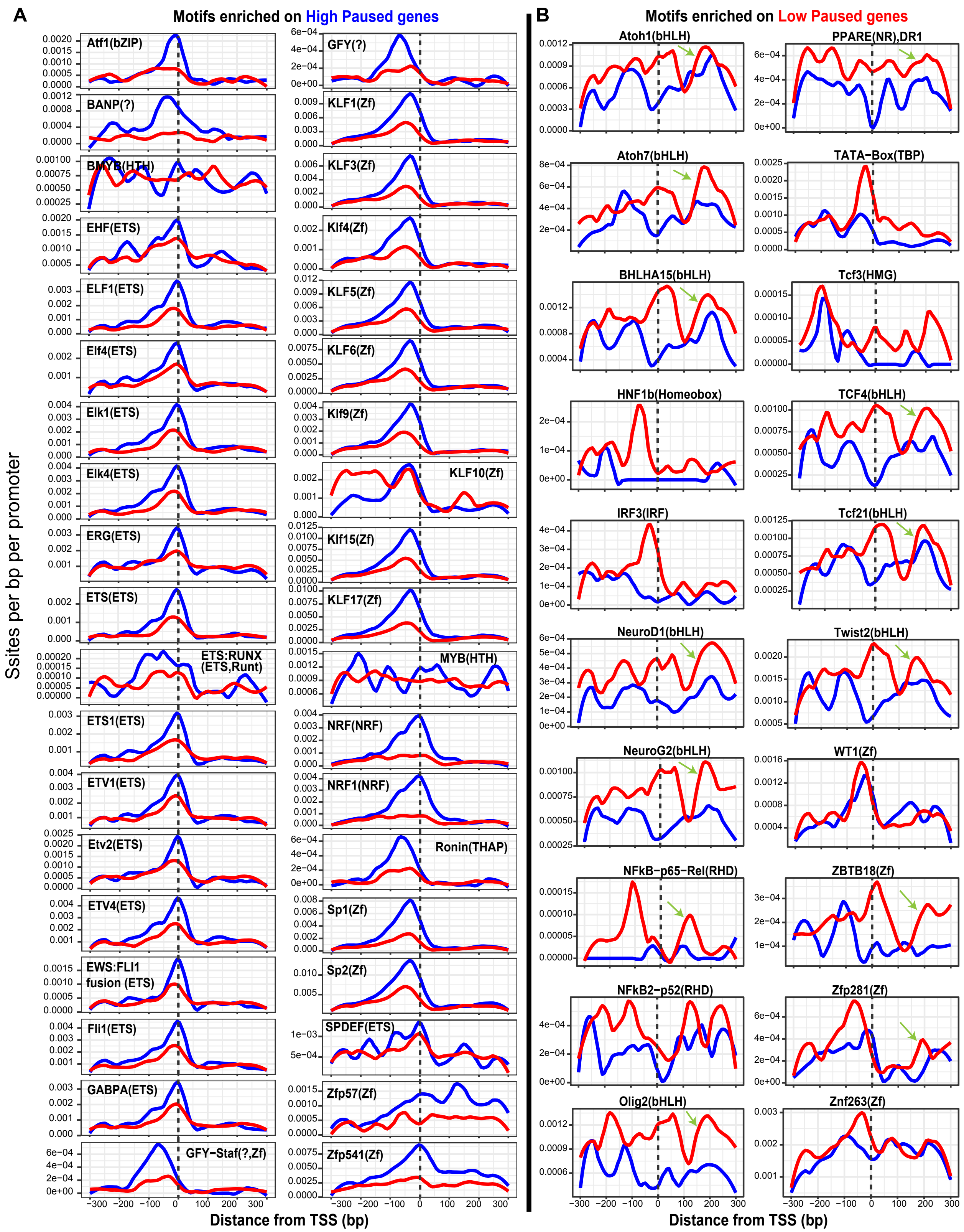
A
Motifs enriched on High Paused genes
B
Motifs enriched on Low Paused genes
Ssites per bp per promoter
Atf1(bZIP)
BANP(?)
BMYB(HTH)
EHF(ETS)
ELF1(ETS)
Elf4(ETS)
Elk1(ETS)
Elk4(ETS)
ERG(ETS)
ETS(ETS)
ETS:RUNX (ETS,Runt)
ETS1(ETS)
ETV1(ETS)
Etv2(ETS)
ETV4(ETS)
EWS:FLI1 fusion (ETS)
Fli1(ETS)
GABPA(ETS)
GFY-Staf(?,Zf)
GFY(?)
KLF1(Zf)
KLF3(Zf)
Klf4(Zf)
KLF5(Zf)
KLF6(Zf)
Klf9(Zf)
KLF10(Zf)
Klf15(Zf)
KLF17(Zf)
MYB(HTH)
NRF(NRF)
NRF1(NRF)
Ronin(THAP)
Sp1(Zf)
Sp2(Zf)
SPDEF(ETS)
Zfp57(Zf)
Zfp541(Zf)
Atoh1(bHLH)
PPARE(NR),DR1
Atoh7(bHLH)
TATA-Box(TBP)
BHLHA15(bHLH)
Tcf3(HMG)
HNF1b(Homeobox)
TCF4(bHLH)
IRF3(IRF)
Tcf21(bHLH)
NeuroD1(bHLH)
Twist2(bHLH)
NeuroG2(bHLH)
WT1(Zf)
NFkB-p65-Rel(RHD)
ZBTB18(Zf)
NFkB2-p52(RHD)
Zfp281(Zf)
Olig2(bHLH)
Znf263(Zf)
Distance from TSS (bp)
Distance from TSS (bp)

**Fig. S8. – Related to Fig. 4 – Motif analysis on low and high paused genes**

**(A-B)** Plots showing location of all significant enriched motifs (q.value < 0.05) on +300bp to -300bp region relative to TSS of high paused genes (Log2PI >6) in (A) and low paused genes (Log2PI <2) in (B).

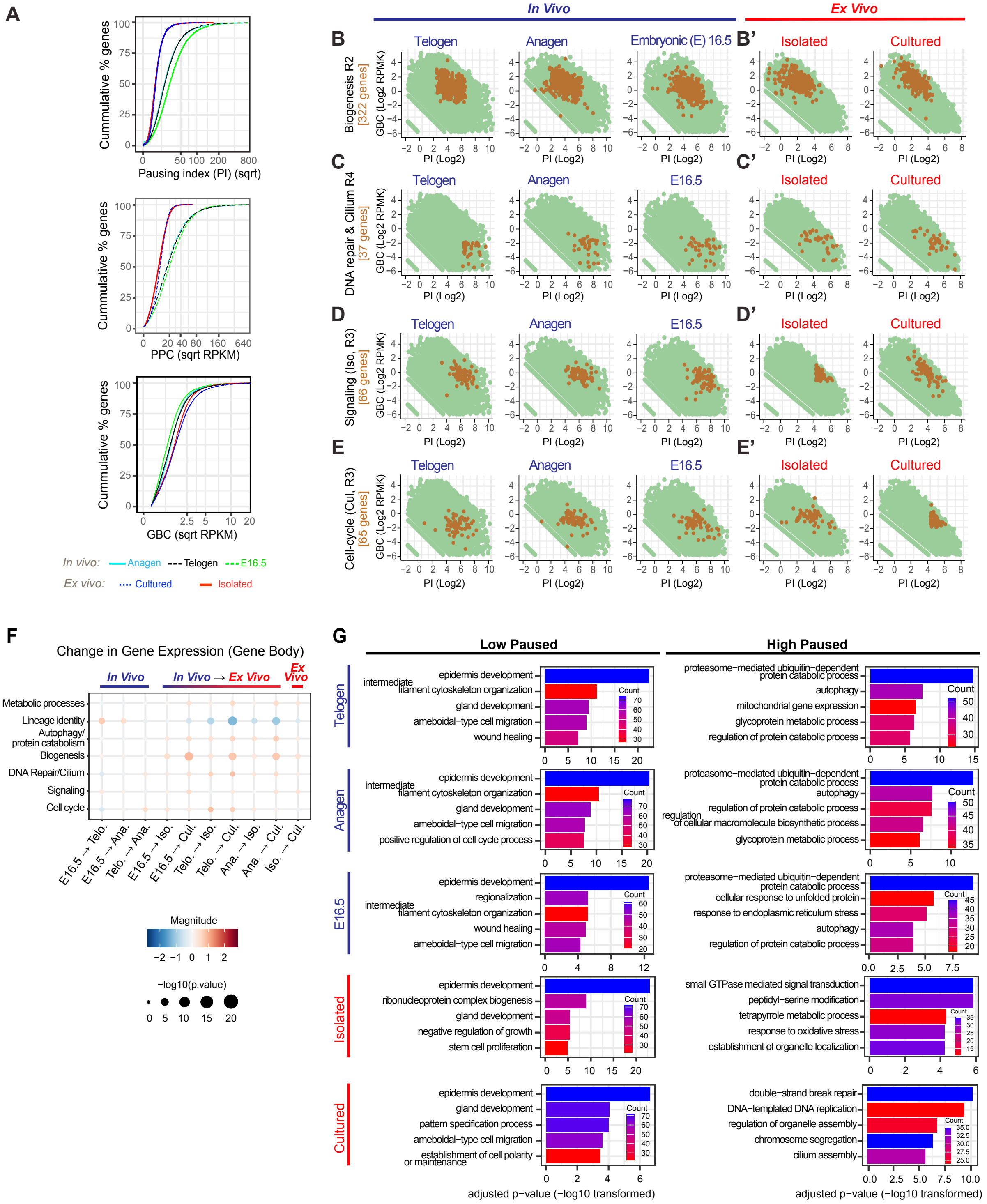

A
Cummulative % genes
Pausing index (PI) (sqrt)
PPC (sqrt RPKM)
GBC (sqrt RPKM)
In vivo: Anagen Telogen E16.5
Ex vivo: Cultured Isolated
In Vivo
Ex Vivo
B
Telogen
Anagen
Embryonic (E) 16.5
B'
Isolated
Cultured
Biogenesis R2 [322 genes]
GBC (Log2 RPMK)
PI (Log2)
C
C'
DNA repair & Cilium R4 [37 genes]
D
D'
Signaling (Iso, R3) [66 genes]
E
E'
Cell-cycle (Cul, R3) [65 genes]
F
Change in Gene Expression (Gene Body)
In Vivo
In Vivo → Ex Vivo
Ex Vivo
Metabolic processes
Lineage identity
Autophagy/ protein catabolism
Biogenesis
DNA Repair/Cilium
Signaling
Cell cycle
E16.5 → Telo.
E16.5 → Ana.
Telo. → Ana.
E16.5 → Iso.
E16.5 → Cul.
Telo. → Iso.
Telo. → Cul.
Ana. → Iso.
Ana. → Cul.
Iso. → Cul.
Magnitude
−log10(p.value)
G
Low Paused
High Paused
Telogen
epidermis development
intermediate filament cytoskeleton organization
gland development
ameboidal−type cell migration
wound healing
proteasome−mediated ubiquitin−dependent protein catabolic process
autophagy
mitochondrial gene expression
glycoprotein metabolic process
regulation of protein catabolic process
Anagen
positive regulation of cell cycle process
regulation of cellular macromolecule biosynthetic process
E16.5
regionalization
cellular response to unfolded protein
response to endoplasmic reticulum stress
Isolated
ribonucleoprotein complex biogenesis
negative regulation of growth
stem cell proliferation
small GTPase mediated signal transduction
peptidyl−serine modification
tetrapyrrole metabolic process
response to oxidative stress
establishment of organelle localization
Cultured
pattern specification process
establishment of cell polarity or maintenance
double−strand break repair
DNA−templated DNA replication
regulation of organelle assembly
chromosome segregation
cilium assembly
Count
adjusted p−value (−log10 transformed)

**Fig. S9. – Related to Fig. 5 – Dynamics of Pol II pausing in various developmental and *ex vivo* conditions**

**(A)** Cumulative distribution plots comparing pausing index (PI), promoter-proximal counts (PPC) and gene body counts (GBC) in keratinocytes of different *in vivo* and *ex vivo* samples. **(B-E')** Scatter plots of GBC vs. PI for different gene groups in both *in vivo* and *ex vivo* conditions. Genes of specific gene groups are colored brown and the remainder are green. (**B-B'**) Biogenesis genes are represented. **(C-C')** DNA repair and cilium genes are represented. **(D-D')** Signaling genes are represented. **(E-E')** Cell-cycle genes are represented. **(F)** Dot plot of the change in GBC of specific gene groups as compared to the rest of the genome across all pair-wise combinations of the five experimental conditions. **(G)** Bar plots of the top unique enriched GO terms for low and high paused genes for *in vivo* and *ex vivo* conditions. (See Materials and Methods for obtaining top unique GO terms).

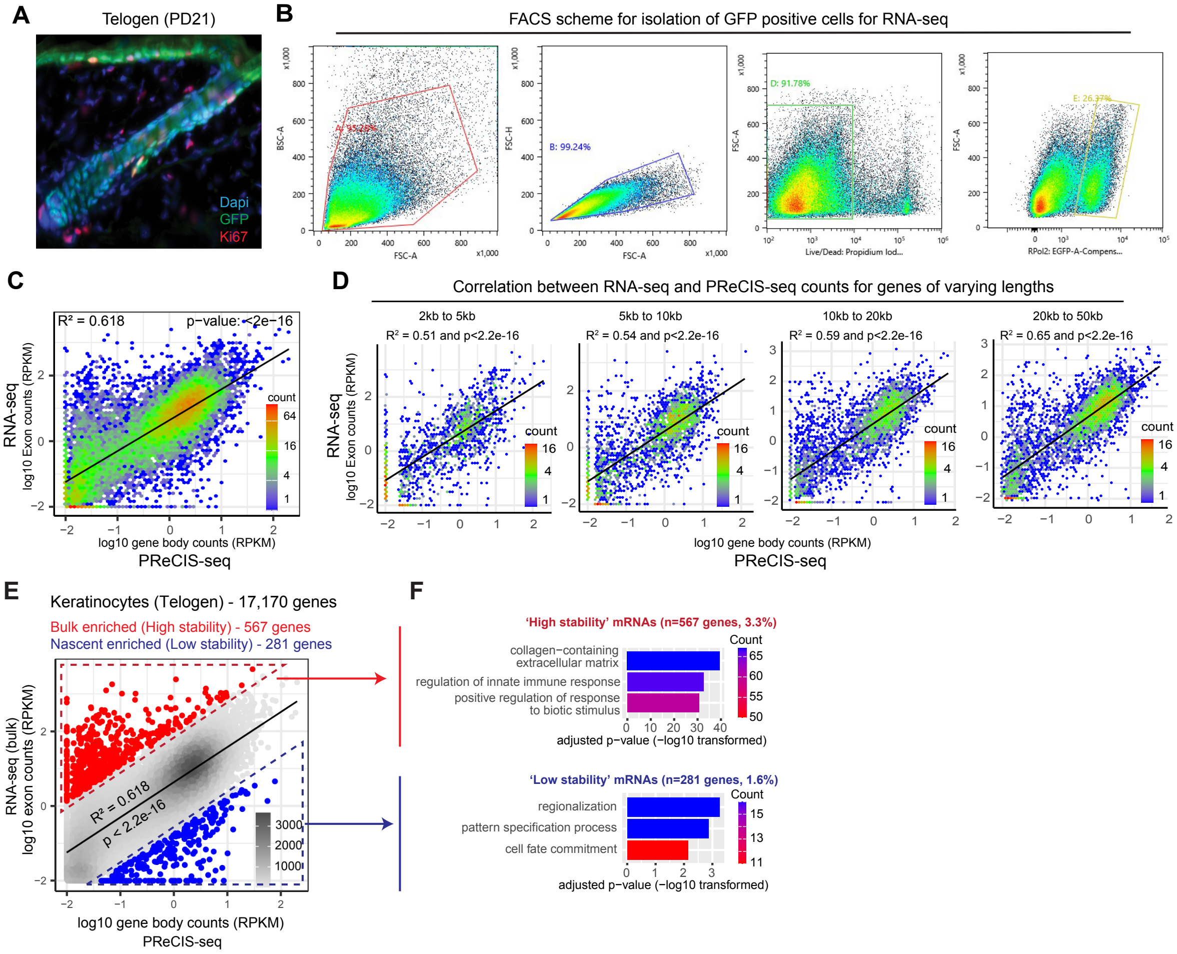


**Fig. S10. Comparisons of nascent RNA levels with steady-state mRNA levels at telogen**

**(A)** Immunofluorescence staining of GFP and Ki67 on skin tissue section from *Polr2b*$^{f/f}$-GFP; K14-Cre mice showing Cre-dependent, nuclear GFP expression specific to keratinocytes in the epidermis. GFP (green), Ki67 (red), DAPI (blue). **(B)** FACS sorting scheme for isolating GFP$^{+}$ cells from *Polr2b*$^{f/f}$-GFP; K14-Cre mice at PD21 for total RNA-seq experiment. (**C**) Comparison of total RPKM-normalized RNA-seq exon counts (y-axis) vs. PReCIS-seq gene body counts (x-axis) for all genes detected in at least one assay (n=17170) at telogen. **(D)** Comparison of total RPKM-normalized RNA-seq exon counts (y-axis) vs. PReCIS-seq gene body counts (x-axis) for genes partitioned into four gene length ranges showing similar correlation across gene lengths different gene length categories. (**E**) Identification of bulk enriched highly stable (>Regression line + 2 Standard Deviation (SD)) and nascent enriched lowly stable (<Regression line – 2 SD) genes by direct comparison of RPKM-normalized RNA-seq exon counts and PReCIS-seq gene body counts. (**F**) Top three enriched GO-terms for highly stable and lowly stable genes at telogen.

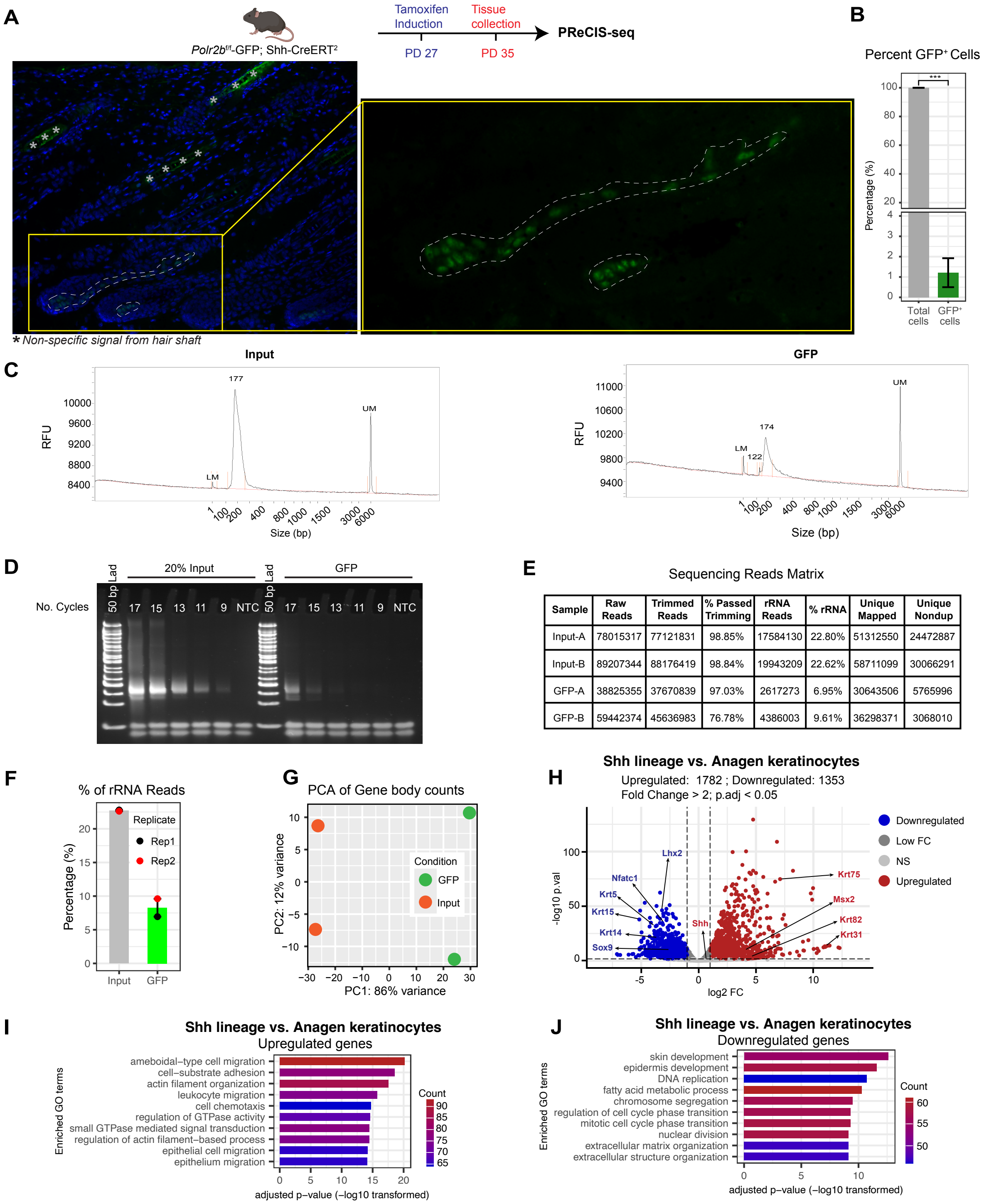


| Sample | Raw Reads | Trimmed Reads | % Passed Trimming | rRNA Reads | % rRNA | Unique Mapped | Unique Nondup |
|---|---|---|---|---|---|---|---|
| Input-A | 78015317 | 77121831 | 98.85% | 17584130 | 22.80% | 51312550 | 24472887 |
| Input-B | 89207344 | 88176419 | 98.84% | 19943209 | 22.62% | 58711099 | 30066291 |
| GFP-A | 38825355 | 37670839 | 97.03% | 2617273 | 6.95% | 30643506 | 5765996 |
| GFP-B | 59442374 | 45636983 | 76.78% | 4386003 | 9.61% | 36298371 | 3068010 |

**Fig. S11. PReCIS-seq profiling of rare hair matrix lineage derived from Shh$^{+}$ cells using *Polr2b*$^{f/f}$-GFP; Shh-CreERT$^{2}$ mice**

**(A-A')** Induction scheme for *Polr2b*$^{f/f}$-GFP; Shh-CreERT$^{2}$ mice to label Shh$^{+}$ cells and its lineage, and fluorescence image of endogenously expressing GFP in Shh$^{+}$ cells and its lineage. Auto-fluorescent hair shafts and artifacts are indicated by asterisks. (**B**) *Polr2b*$^{f/f}$-GFP; Shh-CreERT$^{2}$ mice tamoxifen induced at PD27, and tissue collected at PD35 showed ~1% of GFP$^{+}$ cells in entire skin tissue. GFP$^{+}$ cells were quantified from whole skin (n=16 images at 20x magnification). Two-sample t-test was used to calculate significance. **(C)** Bioanalyzer profile of PReCIS-seq libraries of Input and GFP pull-down samples from back skin of *Polr2b*$^{f/f}$-GFP; Shh-CreERT$^{2}$ mice (RFU - Relative Fluorescence Units). **(D)** PCR of PReCIS-seq libraries showing quality and fragment distribution. **(E)** Table summarizing mapped read counts before and after de-duplication. **(F)** PReCIS-seq data analysis showing depletion of ribosomal RNA (rRNA) reads in GFP pull-down relative to input control. **(G)** PCA plot showing high correlation of biological replicates (two mice) for gene body counts and a clear separation of GFP pulldown samples as compared to input controls. **(H)** Volcano plots showing differentially expressed genes between GFP pull-down from *Polr2b*$^{f/f}$-GFP; Shh-CreERT$^{2}$ mice vs. GFP pull-down from *Polr2b*$^{f/f}$-GFP; K14-Cre mice (at anagen). Hair matrix lineage genes (e.g., *Krt75*, *Msx2*, *Krt82*, *Krt31*) were upregulated while non matrix basal keratinocyte markers were downregulated, as expected (data S19). **(I-J)** Bar plots showing enriched Gene Ontology terms of significantly up- and down-regulated genes in the Shh lineage as compared to anagen keratinocytes, which showed enrichment of migration associated processes in matrix progenitor lineage as they move upward and depletion of skin development and proliferation associated functions, as expected.

**<u>Description of supplementary data files:</u>**

**Data S1. (separate file)**
List of keratinocytes, mesenchymal, endothelial and immune lineage specific genes

**Data S2. (separate file)**
PPC and GBC for all genes in Input, 8WG16 and GFP pulldown at telogen

**Data S3. (separate file)**
PPC and GBC for all genes in Input and GFP pulldown at telogen and anagen

**Data S4. (separate file)**
Differentially expressed genes at anagen by PReCIS-seq

**Data S5. (separate file)**
Summary of motif analysis results for pause-release and Pol II recruitment regulated genes

**Data S6. (separate file)**
List of functional category specific genes

**Data S7. (separate file)**
List of non-lineage genes used for calculating background expression cut off

**Data S8. (separate file)**
GO analysis results for R1 to R6 genes at telogen

**Data S9. (separate file)**
List of all identified dREG elements in telogen keratinocytes

**Data S10. (separate file)**
Overlapping and non-overlapping dREG elements with cCREs from ENCODE database

**Data S11. (separate file)**
List of nearest dREG elements for different gene categories based on expression levels

**Data S12. (separate file)**
List of nearest dREG elements for different gene categories based on PI levels

**Data S13. (separate file)**
List of nearest dREG elements for genes of different GO categories

**Data S14. (separate file)**
Summary of motif analysis of high and low paused genes using HOMER

**Data S15. (separate file)**
PPC and GBC for all genes in E16.5, Isolated and Cultured keratinocytes

**Data S16. (separate file)**
GO analysis results for low and high paused genes for all five conditions

**Data S17. (separate file)**
Quantification of GFP+ cells in back skin of *Polr2b*$^{f/f}$-GFP; Shh-CreERT$^2$ mice

**Data S18. (separate file)**
PPC and GBC for all genes in input and GFP pull-down of *Polr2b*$^{f/f}$-GFP; Shh-CreERT$^2$ mice

**Data S19. (separate file)**
Differentially expressed genes in Shh-CreERT$^2$ targeted cells as compared to K14Cre targeted cells at anagen

**Description of supplementary table:**

**Table S1. (separate file)**
Summary of motif enrichment analysis for enhancers and promoters
